# Modeling Alpha-Synuclein Pathology in a Human Brain-Chip to Assess Blood-Brain Barrier Disruption in Parkinson’s Disease

**DOI:** 10.1101/2020.07.22.207340

**Authors:** Iosif Pediaditakis, Konstantia R. Kodella, Dimitris V. Manatakis, Chris D. Hinojosa, Elias S. Manolakos, Lee L. Rubin, Geraldine A. Hamilton, Katia Karalis

## Abstract

Parkinson’s disease and related synucleinopathies are characterized by the abnormal accumulation of alpha-synuclein aggregates, loss of dopaminergic neurons, and gliosis in the substantia nigra. Although clinical evidence and *in vitro* studies indicate disruption of the Blood-Brain Barrier in Parkinson’s disease, the mechanisms mediating the endothelial dysfunction remain elusive. Lack of relevant models able to recapitulate the order of events driving the development of the disease in humans has been a significant bottleneck in the identification of specific successful druggable targets. Here we leveraged the Organs-on-Chips technology to engineer a human Brain-Chip representative of the substantia nigra area of the brain containing dopaminergic neurons, astrocytes, microglia, pericytes, and microvascular brain endothelial cells, cultured under fluid flow. Our αSyn fibril-induced model was capable of reproducing several key aspects of Parkinson’s disease, including accumulation of phosphorylated αSyn (pSer129-αSyn), mitochondrial impairment, neuroinflammation, and compromised barrier function. This model is poised to enable research into the dynamics of cell-cell interactions in human synucleinopathies and to serve as testing platform for novel therapeutic interventions, including target identification and target validation.

## Introduction

In Parkinson’s Disease (PD) and related synucleinopathies, the accumulation of alpha-synuclein (αSyn) plays a key role in disease pathogenesis. Pathological assessment of post-mortem brains from PD patients has demonstrated abnormal inclusions, enriched in misfolded and aggregated forms of αSyn, including fibrils^1,2^. These findings together with a wealth of experimental data, support the hypothesis for a central role of αSyn aggregation in the formation of the Lewy bodies and, therefore in the pathogenesis of synucleinopathies^3–5^. Recently, αSyn has been identified in body fluids, such as blood and cerebrospinal fluid^6,7^, and has been postulated to be also produced by peripheral tissues^8,9^. However, the ability of αSyn to cross the blood-brain barrier (BBB) in either direction and its potential contribution to the endothelial dysfunction described in patients with PD^10–13^, remains unclear.

Experimental models of PD, such as animal models^14,15^ or conventional cell culture systems^16–18^, have advanced our understanding of the role of αSyn and its aggregated forms in the development of the disease and the neuronal toxicity. However, these models have not been able to uncover the dynamics of the specific interactions between the events in the brain parenchyma and the BBB, either in normal or pathological conditions. Animal models have so far shown a minimal ability for translation of their findings to human patients, and present with limitations to follow the cascade of tissue responses, that require specialized imaging and frequent sampling^19^. Conventional cell culture systems, including co-culture in Transwells, also have limitations such as difficulty to maintain nutrient concentrations, lack of fluid flow, and compromised ability, if any, to recapitulate the cell-cell interactions and cytoarchitecture at the neurovascular unit^20,21^. Recently, microengineered Organs-on-Chips^22,23^ have been successfully developed for multiple complex organs, including intestine, lung, liver, heart, and brain^24–29^. Organs-on-Chips enable the recreation of a more physiological mircroenvironement, including co-culture of cells on tissue-specific extracellular matrices (ECM), the application of flow to enable a dynamic microenvironment and *in vivo*-relevant mechanical forces such as fluidic sheer stress. There have been several approaches to model the Blood-Brain Barrier (BBB) in a chip, with models that have been designed to reconstitute the cerebrovascular interface, however they often have not included combinations of key cell types, such as region-specific neurons, astrocytes, and microglia, to more fully reconstruct the complex physiology of neurovascular unit milieu^30–34^.

In the present study, we describe a novel approach where we have developed a human Brain-Chip with dopaminergic neurons of the substantia nigra, an area predominantly affected in PD (referred to here as the “Substantia Nigra Brain-Chip”). Our Substantia Nigra Brain-Chip recreates the vascular-neuronal interface, and it is populataed with human iPS-derived brain endothelial cells, pericytes, astrocytes, microglia, and dopaminergic neurons. In order to model states of exposure to abnormal αSyn aggregation and confirm the capability of the Substantia Nigra Brain-Chip to generate clinically relevant endpoints, we developed a model of synucleinopathy by introducing human αSyn pre-formed fibrils (PFFs or “aSyn fiblirs”) within the brain channel. We provide evidence that this model can replicate pathological hallmarks observed in human PD brains, including pSer129-αSyn accumulation, mitochondrial dysfunction, and progressive neuronal death^35^. In parallel, we demonstrate activation of astrocytes and microglia, in line with the active inflammatory process operating in the substantia nigra in patients with PD^36^. Further, we provide evidence that the worsening of the brain pathology over time impacts the whole neurovascular unit, as evidenced by compromised BBB permeability.

When taken together, these data suggest that the human αSyn fibril-induced disease model that we have established on the Substantia Nigra Brain-Chip provides a new model for recapitulating and investigating complex pathophysiological features of PD, including the BBB dysfunction. This model can thus be employed as a platform for target identification, target validation, and evaluating the efficacy of new therapies against PD and other synucleinopathies.

## Results

### Development and characterization of a human Substantia Nigra Brain-Chip

To develop the Substantia Nigra Brain-Chip, we leveraged our previously described Organs-Chip design^25^, which consists of two microfluidic channels fabricated from polydimethylsiloxane (PDMS) elastomer separated by a thin (50 μm) PDMS membrane containing multiple pores (7 μm diameter, 40 μm spacing). Each channel has dedicated inlet and outlet ports for the seeding of human cells that are maintained under controlled laminar flow applied independently to each channel (**Fig. 1a**). The membrane separating the two channels is coated on both sides with tissue-specific ECM cocktail, optimized for the Brain-Chip to contain collagen type IV, fibronectin, and laminin. First (D0), we seeded in the brain channel human iPS-derived dopaminergic (DA) neurons derived from a healthy donor, as well as human primary brain astrocytes, microglia, and pericytes at respective seeding densities, as described in the Materials section. The next day (D1), we seeded human iPS-derived brain microvascular endothelial cells (HBMECs) on the opposite surface of the membrane (**Fig. 1b**). Glia (astrocytes and microglia) and pericytes cultured in the brain channel support the proper development and maintenance of the BBB function, as previously reported^33,37,38^. The brain and vascular channels were perfused with endothelial cell medium supplemented with 2% platelet-poor plasma-derived serum, and specific Dopaminergic Neurons Media, respectively (see Methods). The Substantia Nigra Brain-Chip was maintained for two days (D1-D2) in static culture to promote the formation of the endothelial lumen and acclimate cells to the microenvironment before switching to continuous medium flow (60 μL h^-1^). Double-label immunofluorescence with antibodies against tyrosine hydroxylase (TH) and Microtubule-Associated Protein 2 (MAP2) after 8 days in culture, revealed the vast majority of neurons as TH-positive (∼80%), confirming their identity as midbrain dopaminergic neurons (**Fig. 1c**). We also assessed the secreted levels of dopamine via enzyme-linked immunosorbent assay (ELISA) to confirm the functionality of the dopaminergic neurons in the Brain-Chip (**Supplementary Fig. 1a**). Similarly, the other cells of the co-culture in the brain channel of the Substantia Nigra Brain-Chip assessed on D8 of the culture, were found to express the cell-specific markers glial fibrillary acidic protein (GFAP; astrocytes), transmembrane protein 119 (TMEM119; resting microglia), and proteoglycans (NG2; pericytes) (**Supplementary Fig. 1b**). Development of tight junctions in the endothelial monolayer in the vascular channel of the Substantia Nigra Brain-Chip was shown by the expression of Claudin-1, Claudin-5, Occludin, ZO-1 as well as the cell-cell adhesion protein CD31 (**Fig. 1d, e**), as described for the cerebral endothelial cells of the human blood-brain barrier^33,39^. The Substantia Nigra Brain-Chip sustained barrier integrity for up to 7 days in culture under continuous perfusion (D8), as assessed by low passive diffusion of dextran Cascade Blue (Mw: 3 kDa), and lucifer yellow (Mw: 0.5 kDa). Specifically, the apparent permeability of the BBB in the Substantia Nigra Brain-Chip was maintained at values within a range of 1-3×10^−6^ cm s^-1^ and 4-6×10^−6^ cm s^-1^, for dextran (3 kDa) and luciferin yellow (0.5kDa) respectively, evidence of the size-dependent transport across the BBB on Substantia Nigra Brain-Chip (**Fig. 1f**). Notably, the low permeabilities of the Brain-Chip to dextran were comparable to previously reported *in vivo* values^40,41^.

**Figure 1.**
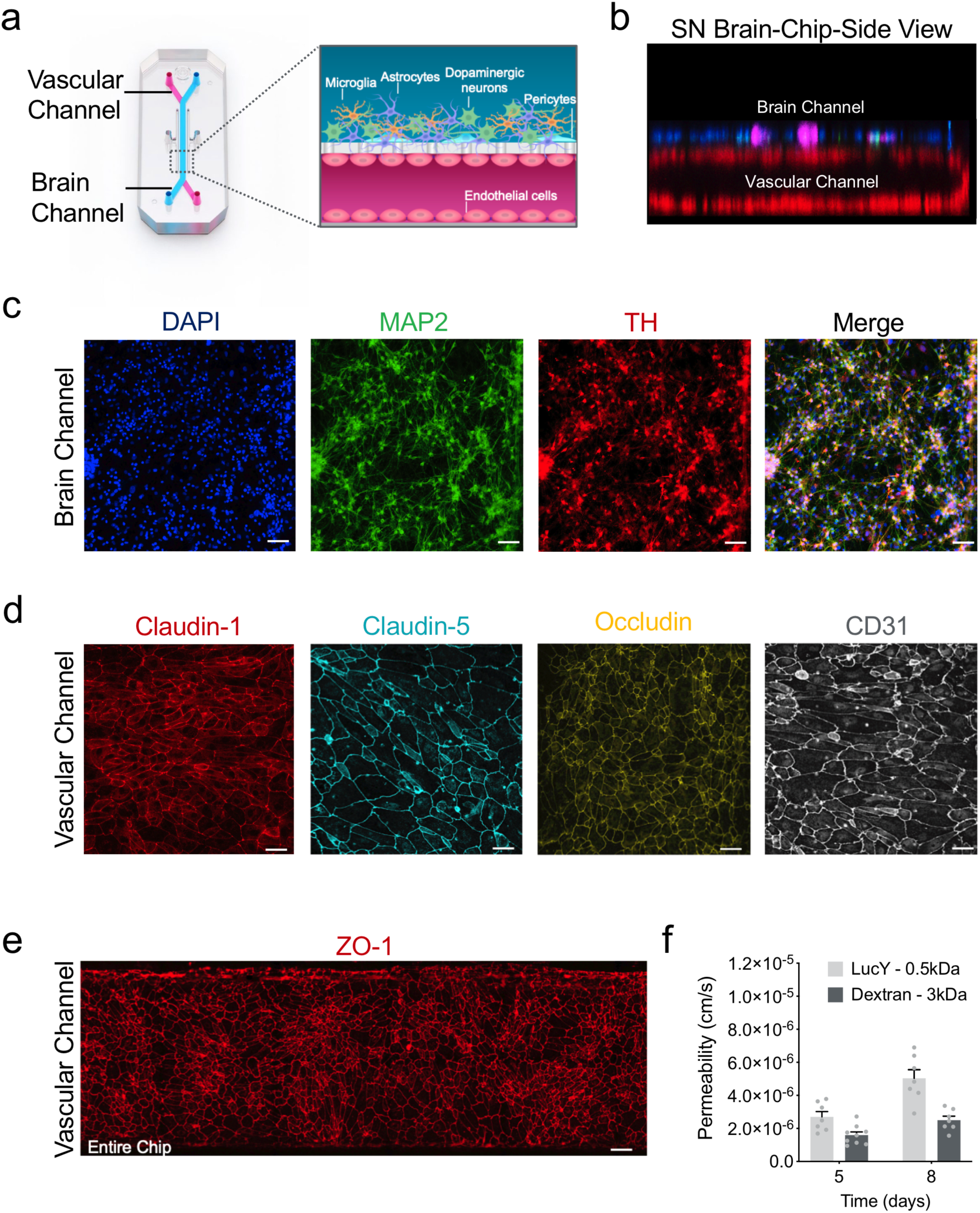
Reconstitution of the human substantia nigra Substantia Nigra Brain-Chip. a) Schematic depiction of the Substantia Nigra Brain-Chip of a two-channel microengineered chip with iPS-derived brain endothelial cells cultured on all surfaces of the lower vascular channel, and iPS-derived dopaminergic neurons, primary human brain astrocytes, microglia and pericytes on the upper surface of the central horizontal membrane in the upper brain channel. b) 3D reconstruction of a confocal z-stack showing the organization of all five cell types in the Substantia Nigra Brain-Chip. c) Representative image of iPS-derived dopaminergic neurons that are stained with DAPI (blue) MAP2 (green), TH (red), and a merged image on day 8. Scale bars: 100 μm. d) Immunofluorescence micrographs of the human brain endothelium cultured on the vascular channel of Brain-Chip for 7 days post-seeding (D8) labeled with Claudin-1 (red), Claudin-5 (cyan), Occludin (yellow), and CD31 (white). Scale bars: 100 μm. e) Immunofluorescence micrographs demonstrate high levels of expression of ZO-1 (red) across the entire endothelial monolayer. Scale bars: 100 μm. f) Quantitative barrier function analysis via permeability to 3 kDa fluorescent dextran, and 0.5 kDa lucifer yellow crossing through the vascular to the neuronal channel on day 5 and 8 (n=6-9 independent chips). Error bars present mean±SEM.

### Transcriptomic profiling of the Substantia Nigra Brain-Chip

Next, we compared the global RNA-sequencing (RNA-seq) profiles data of neurovascular units constructed established conventional cell cultures (n=4), Substantia Nigra Brain-Chips cultured under constant flow (n=4), and human adult brain-derived substantia nigra (n=8) retrieved from the Genotype-Tissue Expression (GTEx) Portal^42^. The conventional cell cultures and the Substantia Nigra Brain-Chips were seeded using the same cell-type composition and subjected to the same experimental conditions (**Fig. 2**).

**Figure 2.**
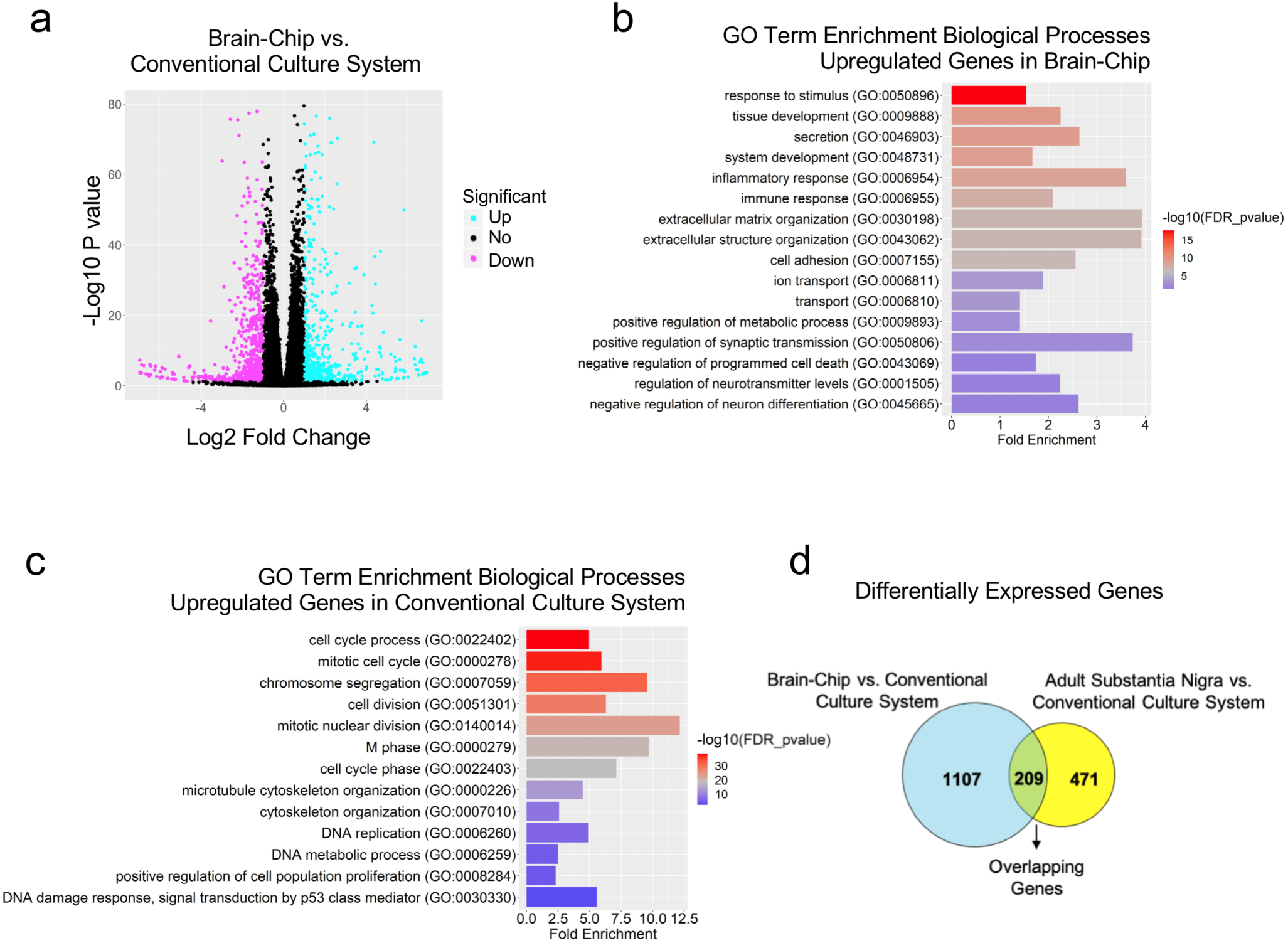
Differentially Expressed (DE) genes and enriched gene ontology categories in Substantia Nigra Brain-Chip or conventional cell cultures as compared to the adult substantia nigra. a) The volcano plot resulted by the DGE analysis between Substantia Nigra Brain-Chip and conventional cell cultures. For the selection of the DE genes we used the following thresholds: adjusted p-value< 0.05 and |Log2(foldchange)| > 1 The identified up- (down-) regulated genes are highlighted in cyan (magenta) color respectively. Sample sizes were as follows: Substantia Nigra Brain-Chip, n=4, conventional cell culture system, n=4. b) and c) List of biological processes identified by Gene Ontology (GO) enrichment analysis using the up- and down-regulated genes respectively resulted by the differentially gene expression analysis between Substantia Nigra Brain-Chip and conventional cell cultures. d) DGE analysis identified up- and down-regulated genes in Substantia Nigra Brain-Chip compared to conventional cell cultures (cyan circle), and human adult substantia nigra compared to conventional cell cultures (yellow circle). Gene lists summarized in the Venn diagram are provided in Extended Data. Sample sizes were as follows: Substantia Nigra Brain-Chip, n=4, Conventional cell culture system, n=4, and adult substantia nigra, n=8 (independent biological specimens). Culture in Brain-Chips and conventional cell cultures were done in parallel. Samples were collected and processed for analyses 8 days post-seeding (D8).

We first performed differential gene expression (DGE) analysis between the Substantia Nigra Brain-Chip and the conventional cell cultures. To select the differentially expressed (DE) genes, we applied the following thresholds: adjusted p-value<0.05 and |log2FoldChange| > 1. Out of the 38,887 genes annotated in the genome, 1316 were significantly differentially expressed, with 646 and 670 genes respectively up- and down-regulated in the Substantia Nigra Brain-Chips (**Fig. 2a**). Then, we performed Gene Ontology analysis utilizing the Gene Ontology knowledgebase, to highlight the biological processes significantly enriched within these gene sets. Among the up-regulated genes in the Substantia Nigra Brain-Chip samples, we identified functional gene sets that significantly clustered under 669 GO terms. These functional gene sets were part of brain-relevant biological processes, including synaptic transmission, ion transport, metabolic and immune processes, extracellular matrix organization, cell adhesion, tissue development, and stimuli-evoked responses (**Fig. 2b**). Compared to the Substantia Nigra Brain-Chip, the transcriptome of the conventional cell cultures was enriched in genes involved in cell division, microtubule cytoskeleton organization implicated in mitosis, and cell cycle processes (**Fig. 2c**). These findings indicate that in the Substantia Nigra Brain-Chip, the cells achieve a more mature and/or differentiated state compared to the cells in the conventional cell cultures, which seems to favor the cell proliferating state. These results are in line with previous studies showing that stem-cell-based tissue models exhibit a higher resemblance to the biological properties of the mature tissue when developed in Organs-on-Chips as compared to conventional cell cultures ^43–45^.

Next, we performed additional DGE analysis, to determine specific gene sets that may underlie the closer similarity between the Substantia Nigra Brain-Chip and the adult substantia nigra tissue, as compared to the conventional cell cultures. For this purpose, we analyzed the differences between the Substantia Nigra Brain-Chip or the adult substantia nigra tissue and the conventional cell cultures (Substantia Nigra Brain-Chip versus conventional cell cultures and adult substantia nigra versus conventional cell cultures). We identified 1316 and 680 DE genes, respectively, from each of the above comparisons, with 209 genes at the intersection of the two (**Fig. 2d**). We reasoned that these 209 DE genes, which were common for the Substantia Nigra Brain-Chip and the human adult substantia nigra tissue versus the conventional cell cultures DE comparisons, would provide insights to biological processes in human substantia nigra tissue that were maintained in the Substantia Nigra Brain-Chip. To get further insights into the biological processes enriched in this gene set, we performed Gene Ontology enrichment analysis. This analysis revealed that the 209 overlapping genes were associated with 25 significant GO terms. Notably, the biological processes enriched in this gene set were associated with essential functions such as secretion, transport, as well as tissue and system development (**Supplementary Fig. 1c**). This data indicates that gene expression patterns characterizing the primary substantia nigra brain tissue are recapitulated by the Substantia Nigra Brain-Chip but not by the conventional cell cultures.

### Establishment of an αSyn fibril model in the Substantia Nigra Brain-Chip

Encouraged by the results of transcriptomic analysis detailed above, we then assessed whether the Substantia Nigra Brain-Chip would respond to abnormal, toxic protein species like those found in synucleinopathies, as reflected in disease-relevant endpoints. To this end, we leveraged the availability of αSyn fibrils, a principal constituent of Lewy bodies shown to exert toxicity in DA neurons^46,47^. First, we characterized the capability of exogenously added fibrils to be accumulated and processed by the cells in the Substantia Nigra Brain-Chip. We added human recombinant αSyn fibrils (4 μg/mL) in the culture medium of the brain channel under continuous flow, on Day 2 of the culture (**Figure 3a)**. Three- and six-days upon-exposure to αSyn fibrils (D5 and D8 of the experiment, respectively), we assessed by immunostaining the abundance of phosphoSer129-alpha-synuclein (phospho-αSyn129), a post-translational modification characteristic of pathogenic αSyn species^48,49^. Our results show that exposure of the Substantia Nigra Brain-Chip to αSyn fibrils was sufficient to induce phosphorylation of αSyn in a time-dependent manner (**Fig. 3b, c, and Supplementary Fig. 2a**). As hypothesized, induction of phospho-αSyn129 was only evident following exposure to αSyn fibrils, as the same amount of αSyn monomer or PBS did not lead to the induction of detectable phospho-αSyn129 in the culture. Immunofluorescence staining verified phospho-αSyn129 accumulation within the TH positive neurons in the Substantia Nigra Brain-Chip (**Fig. 3d**).

**Figure 3.**
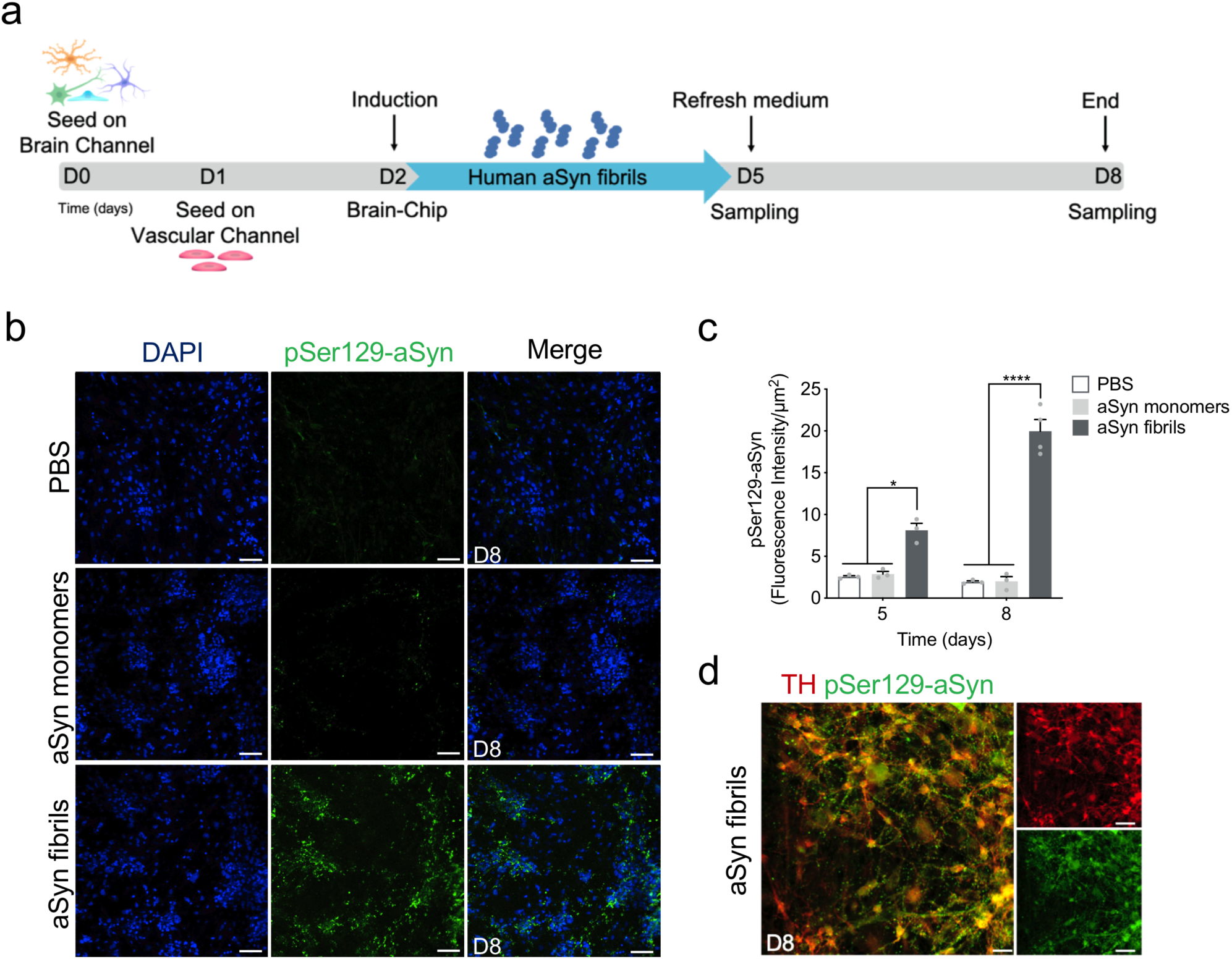
Pathological αSyn accumulation in the brain channel following exposure to human αSyn fibrils. a) Experimental design for assessing the effects of αSyn toxic aggregates (fibrils) in the Substantia Nigra Brain-Chip, including the seeding in the Brain-Chip, the timeline for medium changes, as well as sampling times. b) Immunofluorescence micrographs show the accumulation of phosphorylated αSyn (green, phospho-αSyn129 staining; blue, DAPI) at day six post-exposure (D8). Pathology is absent in the brain channel following exposure to monomer or PBS. Scale bars: 100 μm. c) Quantitative analysis of fluorescence intensity in each group at day three and six post-exposure (D5 and D8, respectively). Statistical analysis is two-way ANOVA with Tukey’s multiple comparisons test (n=3∼4 independent chips with 3∼5 randomly selected different areas per chip, *\*P*<0.05, *\*\*\*\*P*<0.0001 compared to monomeric or PBS group). Error bars present mean±SEM. d) Confocal images are showing double immunostaining for phospho-aSyn129 (green) and TH (red) in the brain channel at six-days post fibrils exposure. Scale bars: 50 μm.

Taken together, our results demonstrate the specific effect of αSyn fibrils on the induction of phospho-αSyn129 pathology in our system.

### Effects of αSyn fibrils in mitochondria and ROS production in the Substantia Nigra Brain-Chip

Emerging evidence indicates the critical role of mitochondrial dysfunction and increased reactive oxygen species in the development of neurodegenerative diseases, including sporadic PD^31,50^. To assess the mitochondrial membrane potential in the cells in the Substantia Nigra Brain-Chip, we used JC-1, a staining probe for the detection of mitochondrial damage. JC-1 in the form of a green monomer enters the cytoplasm and accumulates in the healthy mitochondria, where it forms numerous red J-aggregates. The transition of fluorescence signal from red to green indicates loss of mitochondrial membrane potential, as in cases of significant mitochondrial damage^51^. Exposure to αSyn fibrils led to a lower intensity of red and increased of the green fluorescence, in a time-dependent manner. Reduction in the red-to-green fluorescence intensity ratio was only found in the αSyn fibrils-exposed Substantia Nigra Brain-Chip and not following exposure to the monomeric αSyn species (**Fig. 4a, b, and Supplementary Fig. 2b)**. Next, we measured the intracellular ROS levels in the brain channel of the Substantia Nigra Brain-Chip, on day 8 of the culture, using CellROX reagent. As shown (**Fig. 4c, d**), exposure to αSyn fibrils led to a significant increase in ROS production, as compared to the exposure to αSyn monomers. Thus, we conclude that αSyn fibrils cause functional deficits associated with compromise mitochondrial function in a time dependent manner.

**Figure 4.**
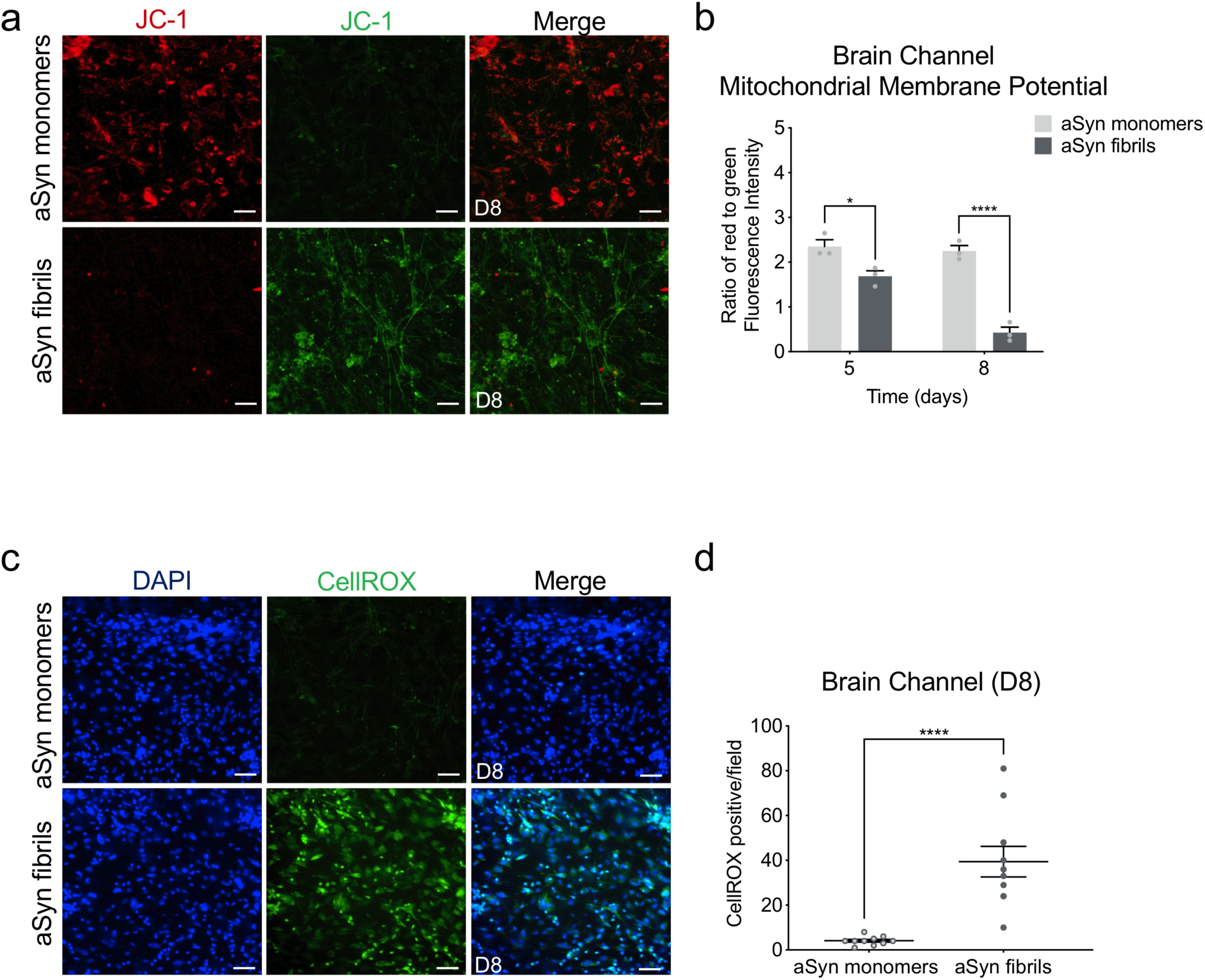
Reduction of mitochondrial activity and increase in ROS production in the αSyn fibril model. a) Mitochondrial membrane potential assessed by JC-1 staining in the brain side at day six post-exposure. Dual emission images (527 and 590nm) represent the signals from monomeric (green) and J-aggregate (red) JC-1 fluorescence. Scale bars: 100 μm. b) Quantitative analysis of the ratio of Red/Green fluorescence intensity in each group at day three and six post-exposure (D5 and D8, respectively). Statistical analysis is two-way ANOVA with Tukey’s multiple comparisons test (n=3 independent chips with 3∼4 randomly selected different areas per chip, *\*P*<0.05, *\*\*\*\*P*<0.0001 compared to monomeric group). c) Representative images of ROS levels (green, CellROX) show higher levels of intercellular ROS in the cells of the brain channel exposed to αSyn fibrils than those exposed to αSyn monomer at day six post-exposure. Scale bars: 100 μm. d) Quantification of the number of CellROX-positive events per field of view in each group. Statistical analysis is Student’s t test (n=3 independent chips with 3∼4 randomly selected different areas per chip, ****p<0.0001 compared to monomeric group). Error bars present mean±SEM.

### αSyn fibrils induce cell death and neuroinflammation in the Substantia Nigra Brain-Chip

Several studies have shown that αSyn fibrils initiate a series of secondary processes leading to neuroinflammation, neurodegeneration, and cell death^46,52^. We first questioned whether the cells in the Substantia Nigra Brain-Chip would respond to αSyn fibrils by induction of apoptosis. Three days (experimental D5) following exposure to αSyn, either monomeric or fibrillar, no effect in cell survival was detected, as reflected by the similar percentage of live cells in all experimental conditions. In contrast, six days upon treatment (experimental D8), there was a significant reduction in live cells in the Substantia Nigra Brain-Chip exposed to αSyn fibrils compare to αSyn monomers or PBS (50.63±3.9 vs 87.02±0.87 vs 91.2±1.05) (**Supplementary Fig. 3a, b**). Confocal immunocytochemical analysis using antibodies against microtubule-associated protein 2 (MAP2) and cleaved caspase-3 (CC3), confirmed the increase in caspase 3-positive neurons on D8 post-exposure to αSyn fibrils, as compared to those exposed to monomeric αSyn (**Fig. 5a, b**). Next, we assessed the extent of inflammatory response mediated by αSyn fibrils in the Substantia Nigra Brain-Chip. We observed increased GFAP staining in the αSyn fibrils-exposed chips suggestive of reactive astrogliosis (**Fig. 5c and Supplementary Fig. 3c**), a known component of the brain inflammatory response^36^. In parallel, there was an activation of microglia, as indicated by the increase in CD68 immunoreactivity (**Fig. 5d and Supplementary Fig. 3d**). As expected, the secreted levels of interleukin-6 (IL-6), and tumor necrosis factor-alpha (TNF-α) induced in the effluent of the neuronal channel, were significantly increased following exposure to αSyn fibrils versus monomeric αSyn (**Fig. 5e, f**). Overall, our data suggest a progressive neurodegeneration linked by strong neuroinflammatory features.

**Figure 5.**
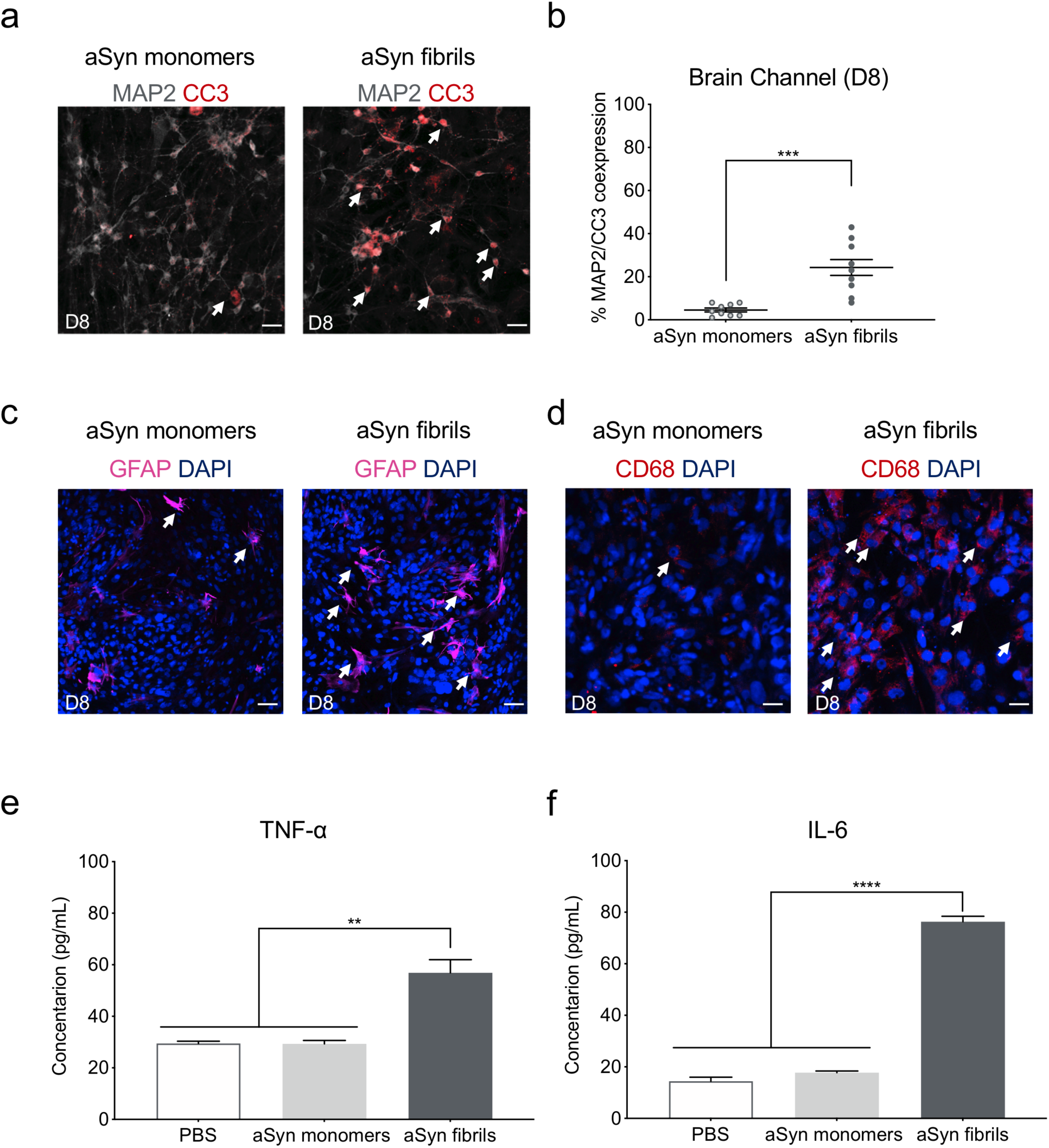
αSyn-induced caspase-3 activation and neuroinflammation. a) Representative merged images showing double immunostaining for MAP2 (grey) and Cleaved Caspase-3 (red, CC3) in the brain channel at six-days post-exposure. Scale bars: 50 μm. b) Quantitative data on the number of CC3 positive neurons. Statistical analysis is Student’s t test (n=3 independent chips with 3∼4 randomly selected different areas per chip, ***p<0.001 compared to monomeric group). c) Immunostaining of the astrocyte marker GFAP (magenta) demonstrated activation of astrocytes at day 8 following exposure to αSyn fibrils compared to monomeric αSyn. Scale bar, 100 μm. d) Immunostaining of the microglial CD68 (red) demonstrated activation of astrocytes and microglia at day 8 following exposure to αSyn fibrils compared to monomeric αSyn. Scale bar, 100 μm. e) The secreted levels of TNF-α in the αSyn fibril model. Statistical analysis was by Student’s t-test (n=6-7 independent chips, **p<0.01). f) The secreted levels of proinflammatory cytokine IL-6 in the αSyn fibril model. Statistical analysis was by Student’s t-test (n=4∼7 independent chips, ****p<0.0001). Error bars present mean±SEM.

### Blood-Brain Barrier disruption in αSyn-associated PD pathology

As the evidence on extraneuronal manifestations of PD is increasing, attention has been drawn to the effects of the disease on the BBB. Measurable levels of αSyn have been identified in the brain and in the systemic circulation, and the current hypothesis is that they are implicated in disease onset and/or progression. The origin of peripheral αSyn remains a subject of discussion, as well as the possibility that αSyn could cross the BBB in either direction^12^. Recent data argues there is BBB dysfunction in PD as in other neurodegenerative diseases and that its role in the pathogenesis and progression of PD may be pivotal^53^. Thus, we ran BBB permeability assays on the Substantia Nigra Brain-Chip upon exposure to αSyn fibrils, as compared to exposure to αSyn monomers or PBS. Our data indicate significantly increased permeability to 160kDa Immunoglobulins (IgG), 3 kDa dextran, and 0.5 kDa lucifer in the brain channel of the Substantia Nigra Brain-Chip six days after exposure to αSyn fibrils (**Fig. 6a, b, and Supplementary Fig. 4a**).

**Figure 6.**
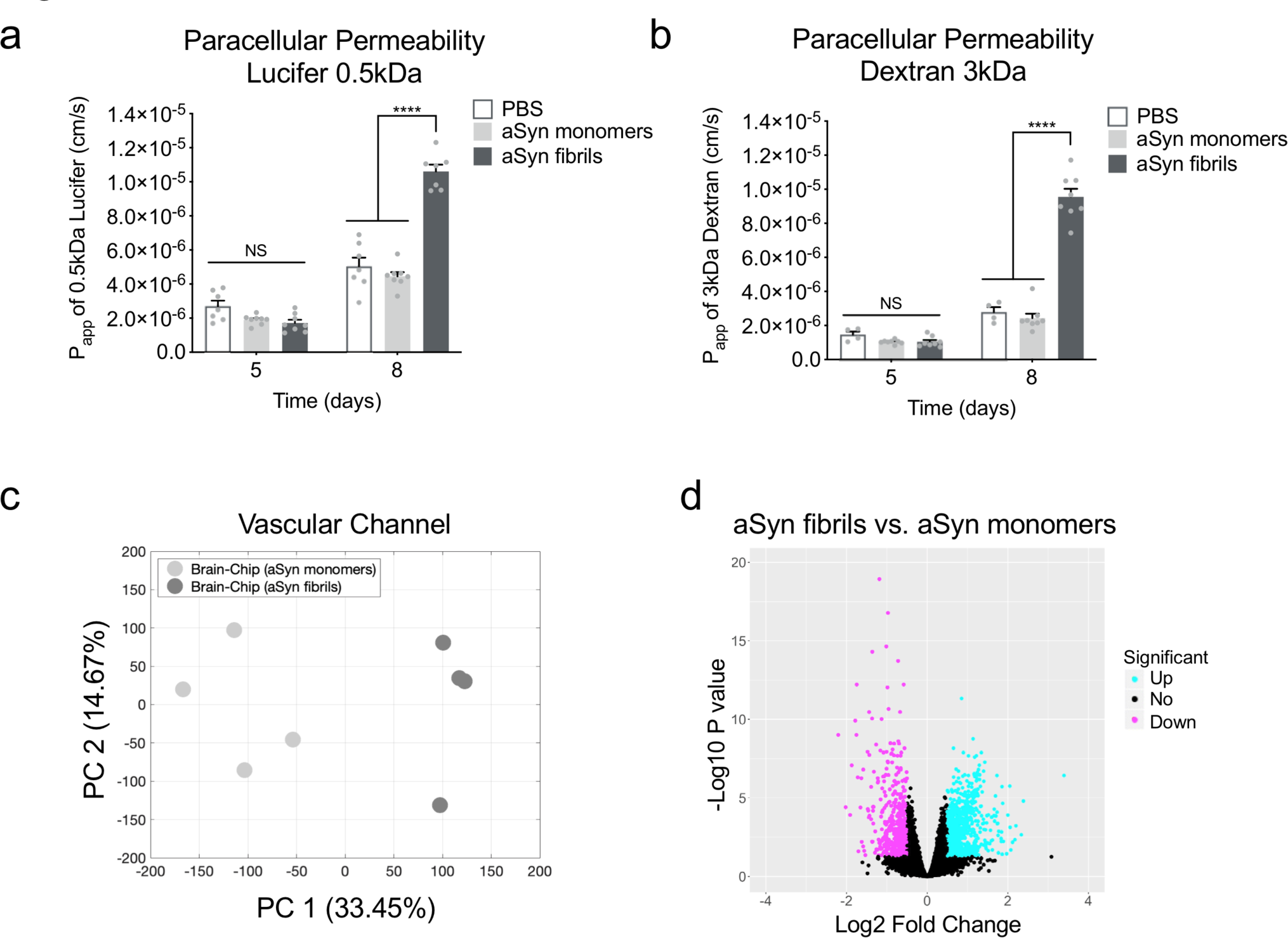
Blood-Brain Barrier dysfunction in the αSyn fibril model. a,b,) Quantitative barrier function analysis via permeability to 0.5 kDa lucifer yellow and 3 kDa fluorescent dextran at day 5 and 8 following exposure to αSyn fibrils or αSyn monomers. Statistical analysis is two-way ANOVA with Tukey’s multiple comparisons test (n=6∼9 independent chips, *\*\*\*\*P*<0.0001 compared to monomeric group, NS: Not Significant). c) Principal component analysis generated using the RNA-seq data generated by the samples collected from the vascular channel of the Substantia Nigra Brain-Chip upon exposure to αSyn monomers or αSyn fibrils (n=4 per condition). A 2D-principal component plot is shown with the first component along the X-axis and the second along the Y-axis. The proportion of explained variance is shown for each component. d) The volcano plot shows the DE genes between αSyn fibrils and αSyn monomers, as identified by the RNA-sequencing analysis. For the selection of the DE genes we used the following thresholds: adjusted p-value< 0.05 and |Log2(foldchange)| > 0.5. The identified up- (down-) regulated genes are highlighted in cyan (magenta) color. Sample sizes were as follows: Brain-Chip (αSyn monomers), n=4, Brain-Chip (αSyn fibrils), n=4.

To further characterize the endothelium in the Substantia Nigra Brain-Chip model and to determine whether the exposure to αSyn fibrils leads to transcriptomic changes in these cells, we ran RNA-seq analysis. Principal components analysis (PCA) revealed differences in the transcriptome profiles between the two conditions, αSyn fibrils and αSyn monomers (**Fig. 6c**). This analysis resulted in the identification of 1280 DE genes, either significantly up-regulated (739 genes) or down-regulated (541 genes) (**Fig. 6d**) in the αSyn fibril-exposed Substantia Nigra Brain-Chips (compete list provided in Supplentary Material). This set of 1280 DE genes includes several genes that have been implicated in BBB dysfunction in a number of diseases^54^. Multiple members of specific gene families were up-regulated, such as extracellular proteases of the Serpin family (SERPINA1), collagens (COL3A1), centromere proteins (CENPE), and kinesins (KIF15). In addition, we identified multiple genes in the αSyn fibrils-exposed chips implicated in key cellular processes associated with PD pathogenesis (**Table 1, Supplementary Fig. 4b)** such as autophagy, oxidative stress, mitochondrial function, inflammation, and vesicular trafficking, highlighting the potential for brain endothelial cells to contribute to molecular mechanisms and functional deficits in PD. Examples of associated genes include leucine-rich repeat kinase 2 (LRRK2)^55^, synphilin-1 (SNCAIP)^56^, monoamine oxidase A (MAOA)^57^, complement 5 (C5)^58^, and apolipoprotein A-1 (APOA1)^59^. Other BBB-related genes with altered expression are low-density lipoprotein receptor-related protein 1B (LRP1B)^60^ and ATP-binding cassette (ABC) transporters (ABCB1)^61^. The upregulation of the LRP1 gene is consistent with previous findings, where dysfunction of LRP1B has been associated with PD^62^. Further, positive and negative associations between specific ABCB1 haplotypes associated with P-glycoprotein activity PD incidence have been reported^61^. Endothelial genes down-regulated upon exposure to αSyn fibrils included the tight junctions claudin gene family (CLDN1, CLDN4, and CLDN9), and the gap junction protein alpha 4 (GJA4). All the above suggest that the Substantia Nigra Brain-Chip could be used to assess novel treatments that protect from vascular dysfunction or improve vascular remodeling in the brain.

**Table 1.** Identification of multiple genes in the αSyn fibrils condition implicated in a variety of cellular processes.

| **Biological functions** | **Genes** | **Regulation** |
| --- | --- | --- |
| Mitochondrial oxidation | MAOA | Up |
| Inflammation | C5, IL1R1, SERPINA1 | Up |
| Autophagy and proteasome system | LRRK2, SNCAIP | Up |
| Vesicular trafficking | CENPE, KIF15 | Up |
| Endothelial active efflux | ABCB1 | Up |
| Lipoprotein receptors | LRP1B, LRP2, APOA1 | Up |
| Solute carrier-mediated transport | SLC16A6, SLC2A6 | Down |
| Tight junctions | CLDN1, CLDN4, and CLDN9 | Down |
| Gap junctions | GJA4 (commonly, Cx37) | Down |

To this purpose, we tested whether the disrupted BBB on the Substantia Nigra Brain-Chip in the αSyn-fibril model could be restored by therapeutic agents targeting mechanisms for clearing of the accumulated αSyn protein. Recent reports of trehalose, a disaccharide approved by FDA, have shown beneficial effects against the accumulation of neurotoxic, aggregated proteins, and neurodegeneration^63^. The study of Hoffmann *et al*. provided evidence that trehalose prevents or halts the propagation of αSyn pathology by targeting lysosomes^64^. Also, studies in aged mice suggest that oral supplementation of the autophagy-stimulating disaccharide trehalose restored vascular autophagy and ameliorated age-related endothelial dysfunction^65^. On the basis of these results, we speculate that trehalose may disturb lysosome integrity and its function, which could subsequently hinder the BBB disruption induced by αSyn fibrils. In this regard, we administered trehalose (10 mM) in the Substantia Nigra Brain-Chip, via the vascular channel on experimental day 5 (three days after adding αSyn fibrils). After 72 hours (experimental D8), we assessed BBB permeability by introducing 3 kDa dextran into the vascular channel (**Fig. 7a**). The trehalose-treated Substantia Nigra Brain-Chips showed significantly decreased BBB permeability (**Fig. 7b**) and rescuing of the damaged tight junctions following exposure to αSyn fibril (**Fig. 7c and Supplementary Fig. 4c)**. Similar resulrs were observed when trehalose was administered in the Substantia Nigra Brain-Chip, via the brain channel (data not shown). Notably, so far, the effects of trehalose have only been evaluated in neuronal cell lines and animal models^64,66,67^. These results suggest that BBB-permeable molecules that reduces protein aggregates and increases BBB integrity might have therapeutic potential for PD and that our model can be used to screen those molecules.

**Figure 7.**
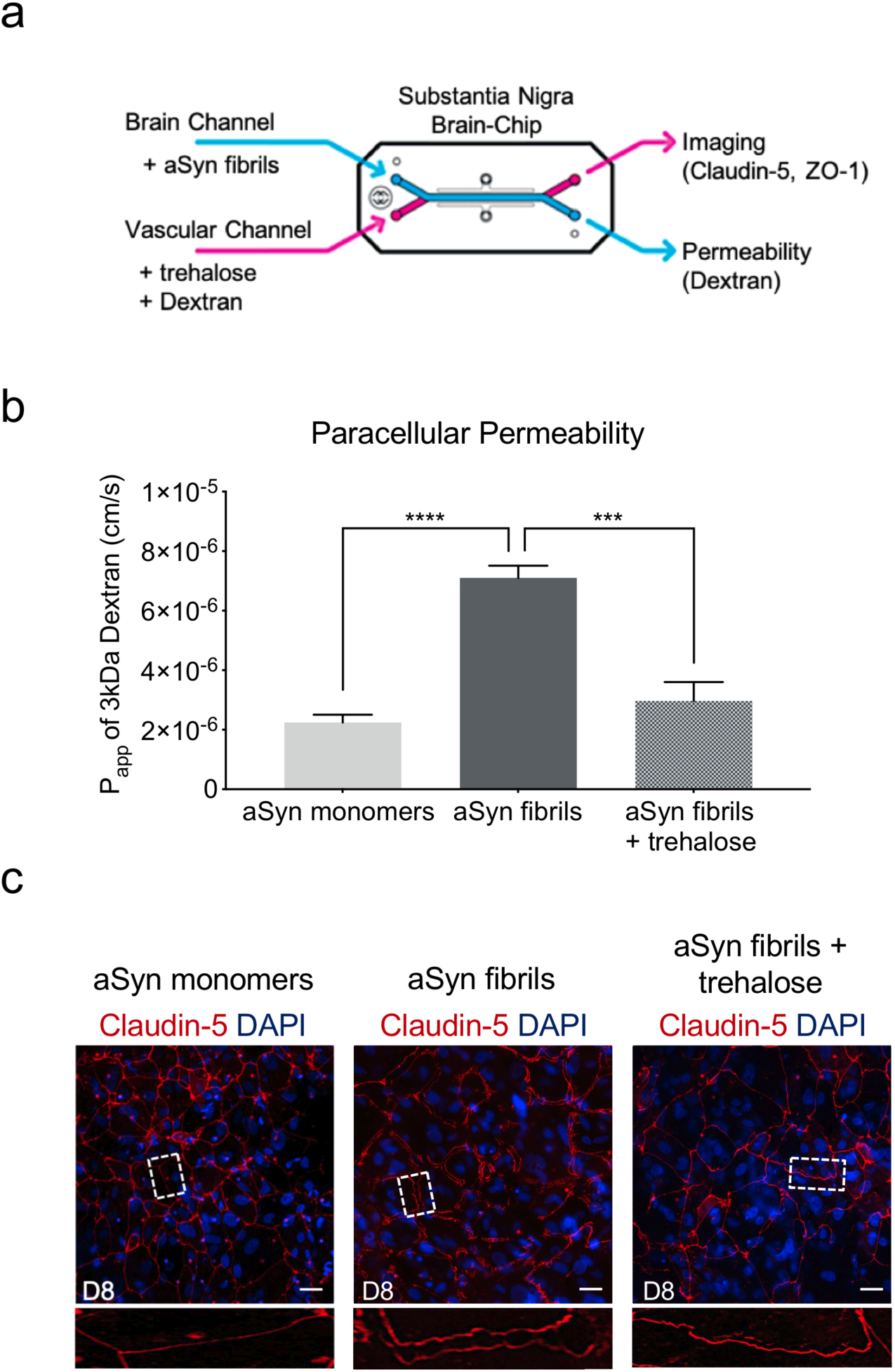
Effect of autophagic inducer, trehalose on BBB integrity. a) Schematic of perfusion of trehalose in the αSyn fibril model. b) Quantitative barrier function analysis via permeability to 3 kDa fluorescent dextran at day 8 in the αSyn fibril model with or without trehalose treatment. Statistical analysis is Student’s t test (n=5∼8 independent chips, *\*\*\*\*P*<0.0001 compared to monomeric group, *\*\*\*P*<0.001 compared to αSyn fibrils). Error bars present mean±SEM. b) Morphological analysis of tight junctions in endothelial cells at day 8 in the αSyn fibril model with or without trehalose treatment. The junction protein expression of Claudin-5 was visualized by immunofluorescence staining with a Claudin-5 antibody and DAPI for cell nuclei. Scale bars: 50 μm.

## Discussion

PD is characterized by an array of premotor and CNS symptoms, together with degenerative changes in the substantia nigra, which expand to more brain areas as the disease advances. Pathology findings reveal the existence of the characteristic Lewy bodies, that are proteinaceous aggregates containing αSyn^1–3^. Experimental models have significantly contributed to our comprehension of the pathogenesis of PD and other synucleinopathies, by demonstrating important aspects of αSyn biology, such as intracellular aggregation and neuronal death^68^. Despite the strong experimental and clinical evidence, the course of events driving the detrimental pathology in synucleinopathies remains unknown, as the diagnosis of the disease usually is made in later stages when the damage has advanced. Furthermore, the existing animal models are limited in their relevance to human disease in terms of disease induction and progression. Given the complexity of the etiology and progress of synucleinopathies and the lack of *in vivo* models representative of the human disease, there is an unmet need for human cell-based models able to recreate the complex biology and uncover the cell-cell interactions driving the tissue pathology.

To address this need, we developed an engineered human Substantia Nigra Brain-Chip to capture the dynamic interactions in the human neurovascular unit, composed of iPSCs-derived brain endothelium and dopaminergic neurons, and primary astrocytes, pericytes, and microglia. A flexible, porus membrane, coated with tissue-relevant ECM, separates the endothelial from the parenchymal cells cultured independently in specific medium, under continuous flow. This system is amenable to imaging and conventional endpoints used in both *in vivo* and *in vitro* studies. The system also enables frequent sampling of the effluent from either side of the membrane for assessment of barrier permeability and the characterization of the secretome at different time points.

Exposure of the Substantia Nigra Brain-Chip to αSyn fibrils led to progressive accumulation of phosphorylated αSyn and the associated induction of specific aspects of αSyn toxicity, such as mitochondrial dysfunction and oxidative stress. Compromised mitochondrial function, as reflected in mitochondrial complex I levels and development of oxidative stress, are central contributors in the neurodegenerative process in PD^50^. We also found that exposure of the Substantia Nigra Brain-Chip to αSyn fibrils, results in microglia activation, astrogliosis, and a time-dependent neuronal loss, as has been described in PD patients^36^. The dynamic, tunable microenvironment of the Substantia Nigra Brain-Chip, which includes physiological relevant mechanical forces and import cytoarchitecture including cell-cell interaction may be the reason we were able to capture complex mechanisms driving key pathologies such as the gradual development of αSyn fibrils-induced toxicity. Another contributing factor might be that the cells on the Brain-Chip transition towards a more mature state, recapitulating aspects of the brain responses in a more physiologically relevant setting that have not been captured by conventional cell culture systems. This hypothesis is supported by our transcriptomic data, and is in agreement with reports showing that maturation of neurons/astrocytes promotes the propensity of αSyn aggregation^47,69^.

The first link between synucleinopathies and inflammation was provided by findings on activated microglia in the substantia nigra of PD patients^70^. Further, marked upregulation of TNF-α and IL-6 mRNA levels was found in the substantia nigra of MPTP-treated animals compared to controls^71^ as well as increased levels of inflammatory mediators in brain tissue from PD patients^72,73^. We similarly detected activated microglia and increased levels of secreted cytokines in the Substantia Nigra Brain-Chip effluent following exposure to αSyn fibrils. Although microglia are understood to be the key driver of the neuroinflammatory responses that propagate the neuronal cell death in PD, additional role(s) for microglia in the progress of synucleinopathies have been suggested^74^. We believe that our Substantia Nigra Brain-Chip provides unprecedented opportunities to identify the exact interactions between microglia and other CNS cell types and how they could be targeted to modify the spread of αSyn pathology. A potential caveat of the current design is the lack of recruited peripheral immune cells, an important component of the disease. However, the perfusion capacity of this platform may be leveraged in the future to model the recruitment of disease-relevant immune cell subsets across the BBB, similar to previous reports^75^.

BBB dysfunction has been recently increasingly viewed as an inherent component of PD progression^10–13^. In PD animal models, including MPTP-treated mice^76^ and 6-hydroxydopamine (6-OHDA)-treated rats^14^, BBB disruption has also been found, in agreement with the clinical data. Additional studies have suggested that αSyn deposition increases BBB permeability^77^ and PD development^78^. Despite the strong experimental and clinical evidence on the BBB disruption in PD, the underlying mechanisms remain unclear, whereas it is suggested that BBB involvement might even precede the dopaminergic neuronal loss in substantial nigra^79^. Finally, recent studies propose a peripheral origin to PD, suggesting the BBB’s involvement in intestine-derived signaling that may induces brain pathology as an early mechanism in PD pathogenesis^80^.

Our results show signs of tight junctions derangement and progressively compromised BBB permeability in response to αSyn fibrils. This is in line with previous studies showing deregulation of claudin as a key determinant of the BBB integrity and paracellular permeability^81^. Transcriptomic analysis of the BBB endothelium from the Substantia Nigra Brain-Chip revealed that αSyn fibrils alter the expression of genes associated with distinct biological processes implicated in PD, including autophagy, oxidative stress, mitochondrial function, and inflammation. Excitingly, control over the amount of αSyn accumulation by treatment with the autophagy inducer trehalose rescued the compromised BBB permeability and the derangement of the tight junctions, suggesting a prospective therapeutic approach for treating compromised BBB implicated in PD.

In conclusion, we report the development of a novel Substantia Nigra Brain-Chip, that upon exposure to αSyn fibrils reproduces *in vivo-*relevant aspects of synucleinopathies. The Substantia Nigra Brain-Chip provides a promising model for the study of the specific disease mechanisms underlying this unmet medical need, including the dynamics of BBB dysfunction. Moreover, the chip may be useful to characterize the response to new PD therapies and identify and evaluate associated biomarkers of the disease.

## Acknowledgments

We would like to thank Dr. Kostas Vekrellis for critical editing. This work was supported by The Michael J. Fox Foundation for Parkinson’s Research (16561; to I.P. and K.K.) and by the National Institute of Health, National Center for Advancing Translational Sciences (UG3TR002188; to C.D.H. and K.K.). The content is solely the responsibility of the authors and does not necessarily represent the official views of the National Institutes of Health. We also thank Brett Clair for scientific illustrations.

## Author Contributions

I.P. designed, performed the experiments, and analyzed the data. K.R.K helped perform experiments and interpret data. D.V.M. processed and analyzed the transcriptomic data, and incorporate the associated data in the manuscript. C.D.H provided insightful input on the engineering aspects of the project. E.S.M. was involved in the bioinformatic analysis and provided comments in the paper. G.A.H. provided insightful comments on the findings through the project and reviewed the manuscript. L.L.R. provided critical feedback and reviewed the manuscript. K.K. and I.P. initiated, supervised the study, and wrote the paper with input from all others.

## Figure legends

**Supplementary Figure 1.**
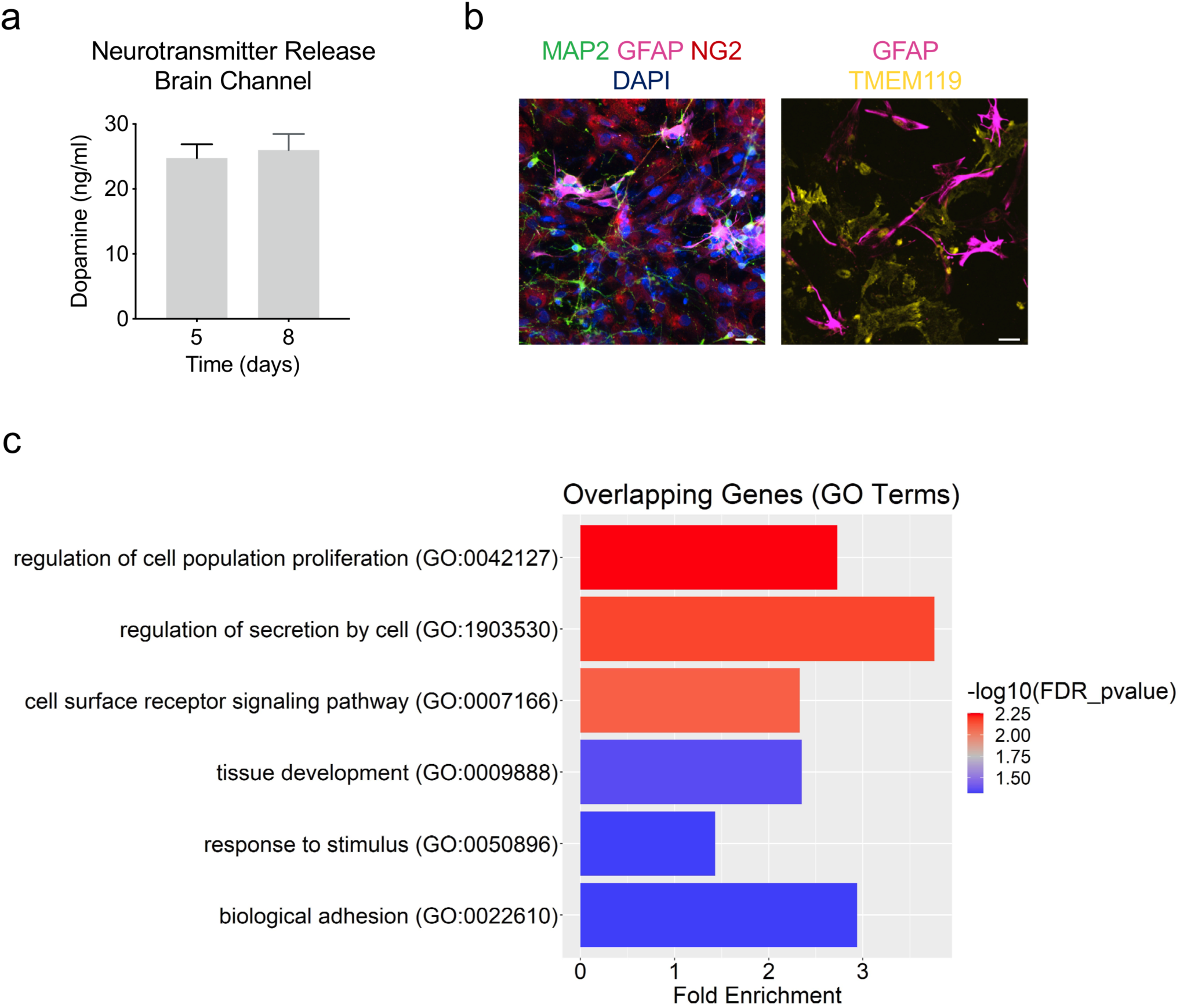
Characterization of the human Substantia Nigra Brain-Chip. a) ELISA for dopamine secreted into the medium of the brain channel on days 5 and 8. (n=3 independent chips with duplicate technical replicates assayed per condition). Error bars present mean±SEM. b) Immunofluorescent microphotographs (left) validate the dopaminergic neurons with MAP2 (green), astrocytes with GFAP (magenta) and pericytes (red), and the DAPI (blue) for cell nuclei. Immunofluorescent microphotographs (right) validate the glia culture: astrocytes (magenta, GFAP staining), and resting microglia (yellow, TMEM119). Scale bars: 50 μm. c) Substantia Nigra Brain-Chip exhibits higher transcriptomic similarity to adult substantia nigra than conventional cell culture. The results of the GO terms analysis using the 209 DE genes showed 6 significantly enriched (FDR adjusted p-value<0.05) biological processes related to tissue development, response to a stimulus, biological adhesion, and cell surface receptor signaling pathway. The size of the bars indicates the fold-enrichment of the corresponding pathways.

**Supplementary Figure 2.**
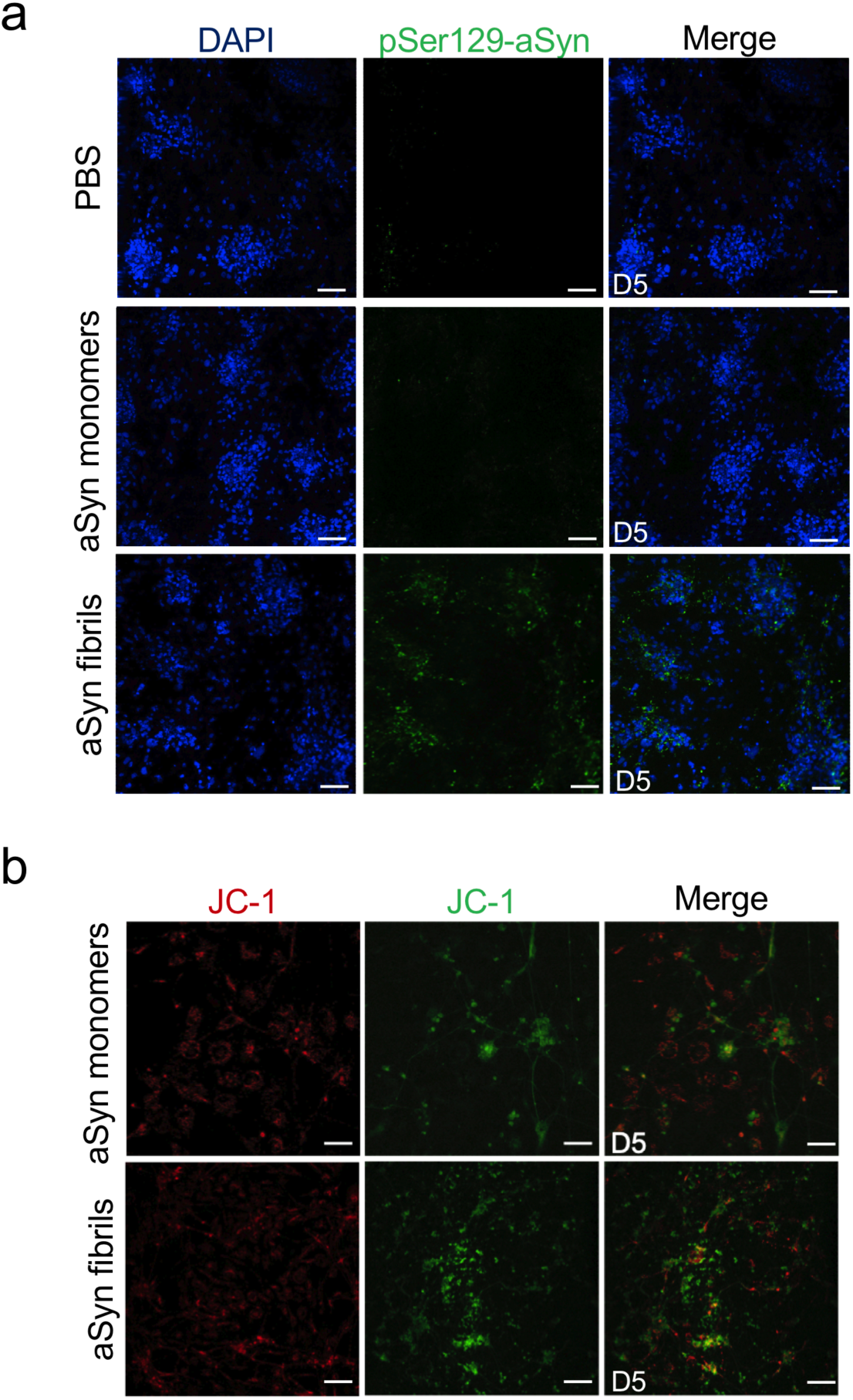
Accumulation of phosphorylated αSyn and mitochondrial impairment in the αSyn fibril model at day 5. a) Assessment of phosphorylated αSyn at three days post-exposure. Immunofluorescence micrographs show the accumulation of phosphorylated αSyn (green, phospho-αSyn129 staining; blue, DAPI). Scale bars: 100 μm. b) Effect of αSyn fibrils on the Mitochondrial membrane potential at three days of posts exposure. Mitochondrial membrane potential assessed by JC-1 staining on the brain side. Dual emission images (527 and 590nm) represent the signals from monomeric (green) and J-aggregate (red) JC-1 fluorescence. Scale bars: 100 μm.

**Supplementary Figure 3.**
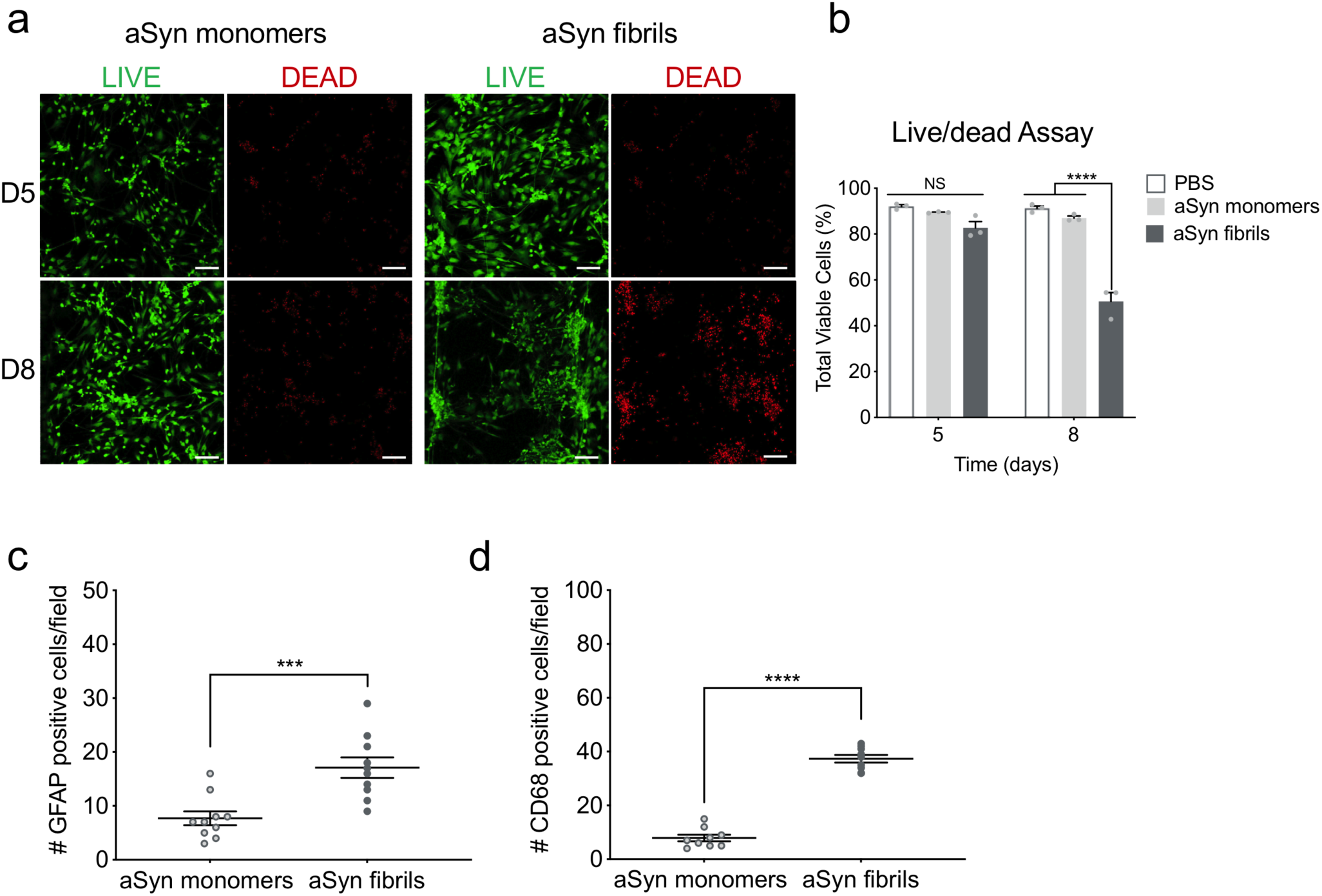
αSyn-induced cell death and neuroinflammation. a) Cell viability (live/dead) assay following exposure to human αSyn fibrils. Live/Dead cell staining assay was designed to test the potential cytotoxicity of αSyn fibrils at days 5 and 8 of culture. Scale bars: 100 μm. b) Data are expressed as the average live cells/total number of cells (sum of calcein AM positive and ethidium homodimer positive cells). Statistical analysis is two-way ANOVA with Tukey’s multiple comparisons test (n=3 independent chips with 3∼5 randomly selected different areas per chip, *\*\*\*\*P*<0.0001 compared to monomeric or PBS group, NS: Not Significant). c) Quantification of the number of GFAP-positive events per field of view. Statistical analysis is Student’s t test (n=3 independent chips with 3∼4 randomly selected different areas per chip, ***p<0.001 compared to monomeric group). d) Quantification of the number of CD68-positive events per field of view. (n=3 Brain-Chips with 3∼5 randomly selected different areas per chip, ****P<0.0001 compared to the monomeric group). Error bars present mean±SEM.

**Supplementary Figure 4.**
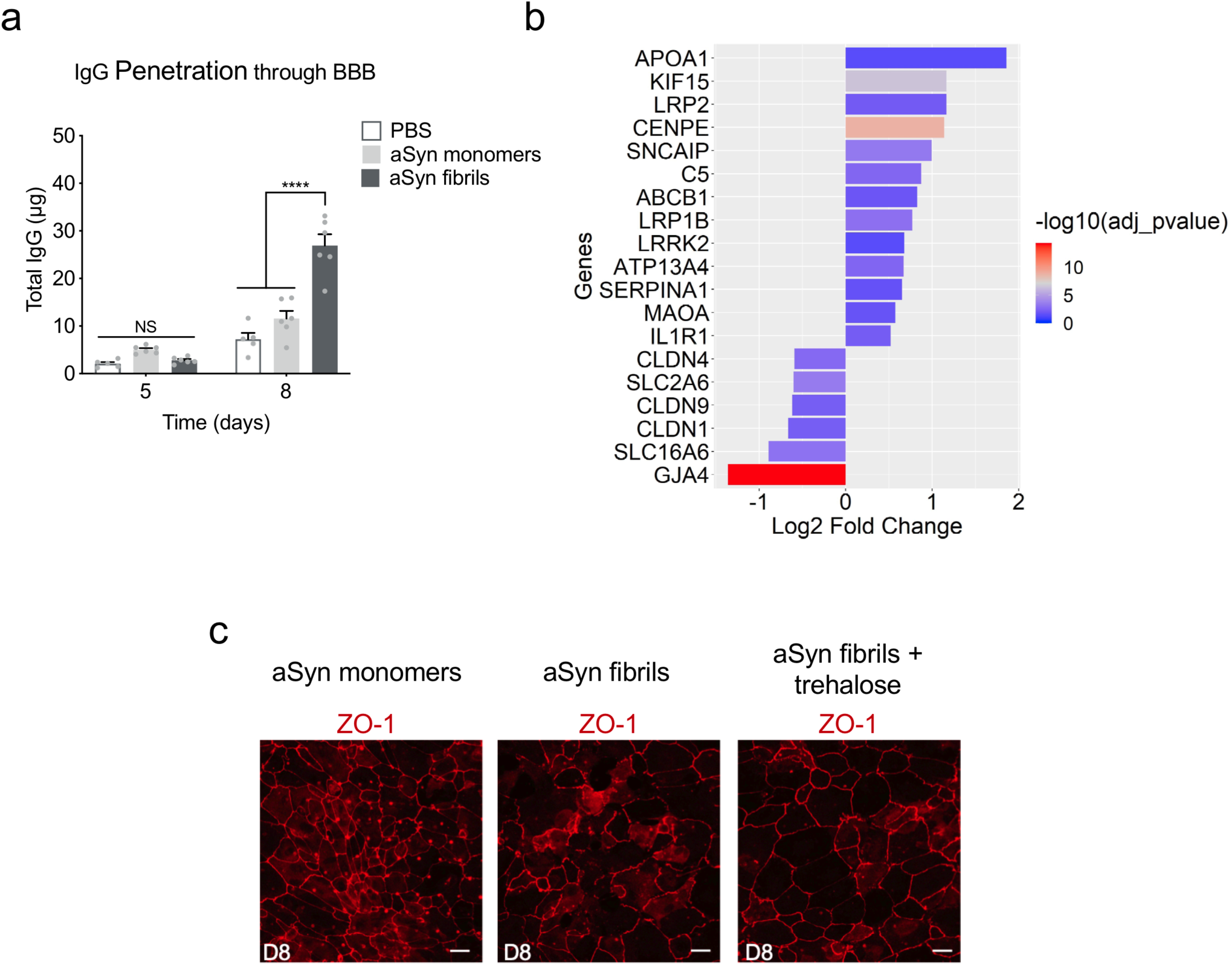
Blood-Brain Barrier dysfunction in the αSyn fibril model. a) Quantitative barrier function analysis via permeability to immunoglobulin G (IgG1) at day 5 and 8 following exposure to αSyn fibrils, αSyn monomers or PBS. Statistical analysis is two-way ANOVA with Tukey’s multiple comparisons test (n=5∼8 independent chips, *\*\*\*\*P*<0.0001 compared to monomeric group, NS: Not Significant). Error bars present mean±SEM. b) A selection from the 739 up-regulated and 541 down-regulated genes identified after performing DGE analysis between αSyn fibrils and αSyn monomers. The size of the bars indicates the log_2_ (Fold-Change) of the corresponding gene expressions and the colors the statistical significance (FDR adjusted p-values) of the corresponding changes. c) Morphological analysis of tight junctions in endothelial cells in the αSyn fibril model with or without trehalose treatment. The junction protein expression of ZO-1 was visualized by immunofluorescence staining with a ZO-1 antibody. Scale bars: 50 μm.

## Methods

### Cell culture

Commercial human iPSC-derived dopaminergic neurons (iCell® Neurons), were purchased from Cellular Dynamics International (CDI, Madison, WI) and maintained in complete maintenance media (iCell DopaNeurons Media). The cells have been characterized by CDI to represent a pure neuronal population with >80% pure midbrain dopaminergic neurons. Primary human astrocytes isolated from the cerebral cortex were obtained from ScienCell and maintained in Astrocyte medium (ScienCell). Primary human brain pericytes were also obtained from ScienCell and maintained in the Pericyte medium (ScienCell). Resting primary human brain microglia were purchased from ATCC and cultured according to the manufacturer’s instructions. The primary cells were used at passage 2-4.

### Brain microvascular endothelial cell differentiation of hiPSCs

Human-induced pluripotent stem cells (hiPSCs) obtained from the Rutgers University Cell and DNA Repository (ND50028; RUCDR) were maintained on Matrigel-coated tissue-culture treated six-well culture plates (Corning) in mTeSR™1 (Stem Cell Technologies). The established hiPSC colonies displayed a normal karyotype in culture. For each independent experiment, we used the same cell passage (P49). Prior to differentiation, hiPSCs were singularized using Accutase^®^ (Invitrogen) and plated onto Matrigel^®^-coated six-well culture plates in mTeSR™1 supplemented with 10 μM Rho-associated protein kinase inhibitor Y27632 (ROCK inhibitor; Stem Cell Technologies) at a density between 25,000 and 50,000 cells cm-2. Directed differentiation of hiPSCs was adapted from a previously reported protocol^38^. Briefly, singularized hiPSCs were expanded for three days in mTeSR™1, then were treated with 6µM CHIR99021 (Stem Cell Technologies) in DeSR1: DMEM/F-12 (Life Technologies), 1% non-essential amino acids (Thermo Fisher Scientific), 0.5% GlutaMAX™ (Thermo Fisher Scientific), 0.1 mM beta-mercaptoethanol (Sigma) to initiate differentiation (day zero). After one day, the medium was changed to DeSR2: DeSR1 plus 1 × B27 (Thermo Fisher Scientific) and changed daily for five days. On day six, the medium was switched to hECSR1: hESFM (Thermo Fisher Scientific) supplemented with 20 ng/mL bFGF (R&D Systems), 10 μM all-trans retinoic acid (Sigma) and 1 × B27. The medium was not changed for 48 hrs. On Day 9, the medium was switched to hESCR2: hECSR1 lacking RA and bFGF. On day ten, cells were dissociated with TrypLE™ (Thermo Fisher Scientific) and replated onto a human placenta-derived collagen IV/human plasma-derived fibronectin/human placenta-derived laminin-coated flasks. After 20 mins, the flasks were rinsed using a medium composed of human serum-free endothelial medium supplemented with 2% platelet-poor plasma-derived serum and 10µM Y27632, as a selection step to remove any undifferentiated cells. Human brain microvascular endothelial cells (HBMECs) were then left in the same medium overnight to allow cell attachment and growth before seeded into the Organ-Chips.

### Brain-Chip microfabrication and Zoë^®^ culture module

Organ-Chips (Chip-S1^®^, Emulate, Inc. Boston, MA, USA) were used to recreate the human Brain-Chip. Chip-S1^®^ is made of a poly(dimethylsiloxane) (PDMS) flexible elastomer. It consists of two channels (1 × 1 mm and 1 × 0.2 mm, “Brain” and “Vascular” channel, respectively) separated by a porous flexible PDMS membrane^25^. Flow can be introduced to each channel independently to continuously provide essential nutrients to the cells, while effluent containing any secretion/waste components from cells is excreted/collected on the outlet of each channel separately. This allows for channel-specific and independent analysis and interpretation of results. The Zoë^®^ culture module is the instrumentation designed to automate the maintenance of these chips in a controlled and robust manner (Emulate, Inc.).

### Human Brain-Chip and cell seeding

Prior to cell seeding, chips were functionalized using Emulate’s proprietary protocols and reagents. After surface functionalization, both channels of the human Brain-Chip were coated with collagen IV (400 μg/mL), fibronectin (100 μg/mL), and laminin (20 μg/mL) overnight-both channels of the chip and then with DopaNeurons medium before seeding cells. A mixture of dopaminergic neurons, astrocytes, microglia, and pericytes was seeded in the upper brain channel of the Brain-Chips at the following concentrations: 2 million cells/mL for dopaminergic neurons, 2 million cells/mL for astrocytes, 0.1 million cells/mL for microglia, and 0.1 million cells/mL for pericytes. After cell seeding, the upper channel of the Brain-Chip was maintained in DopaNeurons medium and incubated overnight at 37°C (Day 0). The following day (Day 1), the lower vascular channel was rinsed with human serum-free endothelial medium supplemented with 2% platelet-poor plasma-derived serum, 10 μM Y27632, and then BMECs were seeded at a concentration of 16-20 million cells/mL to ensure the very tight endothelial monolayer found in the human blood-brain barrier, and the chips were flipped immediately to allow BMECs to adhere to the ECM-coated part of the membrane. After 2 h incubation, the chips were flipped back to let the rest of BMECs sit on the bottom and sides of the channel to form a capillary lumen. The vascular channel of the Brain-Chip was maintained overnight. On Day 2, the Brain-Chips were connected to the Zoë^®^ culture module and perfused continuously through the brain and vascular channel at a flow rate of 30 μl hr^-1^ and 60 μl hr^-1^ respectively, using each channels’ respective media.

### Addition of exogenous alpha-synuclein to brain channel

Human recombinant αSyn monomers and pre-formed fibrils were purchased from Abcam (Monomers; ab218819, Fibrils; ab218819), and were diluted in DopaNeurons Media to a final concentration of 4 μg/mL. On the day of use, αSyn fibrils were sonicated, and their activity was verified by Thioflavin T assay. Endotoxin levels were evaluated by the Limulus amebocyte lysate assay (Endotoxin Testing Services, Lonza Europe), and the amount expressed was negligible. For treatment, freshly prepared monomers and fibrils were used. On Day 2, the upper channel of the Brain-Chip was exposed to monomeric or fibrillar αSyn. After three days of exposure (D5), the medium was changed, and the culture was maintained using DopaNeurons Media (αSyn free) for three more days (D8). Effluents, lysates, and staining were collected/fixed at day three- and day six post-exposure (D5 and D8 respectively), and were analyzed by a microplate reader, ELISA kits, and immunofluorescence microscopy.

### Permeability Assays

Apparent permeability (Papp) of the barrier was calculated by following a previously described method ^82^. Briefly, 100 μg mL^-1^ of dextran (3 kDa) and 20 μg mL^-1^ of lucifer (0.5 kDa) tracers were dosed through the vascular channels for 24 hrs, and concentration of the dextran and lucifer tracers in the outlet samples from both vascular and brain channels was determined by using BioTek (BioTek Instruments, Inc., Winooski, VT, USA). Then, the following equation was used to calculate Papp:

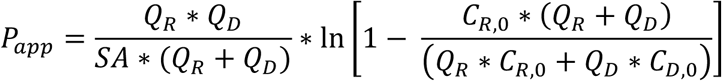

Here, SA is the surface area of sections of the channels that overlap (0.17cm2), QR and QD are the fluid flow rates in the dosing and receiving channels respectively, in units of cm3/s, CR,O and CD,O are the recovered concentrations in the dosing and receiving channels respectively, in any consistent units. IgG permeability was also evaluated after dosing the vascular channel and measuring the IgG content on the brain channel. Detection and quantitation of serum immunoglobulin G (IgG1; Abcam) was performed using the ELISA kit (Abcam), after 24 hrs of perfusion.

### Immunofluorescence microscopy

Chips were fixed with 4% paraformaldehyde in PBS for 10 min and then washed with PBS. Immunostaining was performed after permeabilization in PBS with 0.1% Saponin and blocking for 30 min in 10% donkey serum in PBS with 0.1% Saponin. Immunostaining was performed with specific primary antibodies (anti-TH, anti-GFAP, anti-NG2, anti-TMEM119, anti-pSer129-αSyn, anti-Cleaved Caspase-3, anti-CD68; Abcam), (anti-MAP2, and anti-CD31; Thermo Fisher Scientific), (anti-Claudin-1, anti-Claudin-5, anti-Occludin, anti-ZO-1; Invitrogen), in a 1:100 dilution in 10% donkey serum in PBS with 0.1% Saponin and incubated overnight on the Brain-Chip at 4°C. Fluorescently conjugated secondary antibodies with Alexa Fluor-488, Alexa Fluor-568, or Alexa Fluor-647 were then used when the primary antibodies are not conjugated, and cells were counterstained with DAPI nuclear stain. Images were acquired with either an Olympus fluorescence microscope (IX83) or Zeiss confocal microscope (AxiovertZ1 LSM880).

### Mitochondrial membrane potentials assay

JC-1 probe was employed to evaluate the mitochondrial depolarization in cells seeded at the brain channel. Briefly, cells were incubated with 2 μM of JC-1 dye at 37 °C for 20 min and rinsed twice with PBS, then replaced in fresh medium. Finally, images were taken in the green and red fluorescence channel by confocal laser scanning microscopy imaging. The images were obtained at 488 nm excitation and 530 nm emission for green (JC-1 monomers) and 543 nm excitation and 590 nm emission for red fluorescence (JC-1 aggregates). Four frames per chip at 10X magnification were selected for each treatment, and fluorescence intensity was measured using Fiji/ImageJ.

### Intracellular ROS Measurement

Intracellular ROS production was measured using CellROX^®^ Green Reagent (Thermo Fisher Scientific) according to the manufacturer’s protocol. At day 8, CellROX^®^ reagent was added to the brain channel at a final concentration of 5 μM, and cells were incubated for 60 min at 37 °C in the dark, followed by triple washing with prewarmed PBS. Then, cells were examined with a confocal laser scanning microscope at an excitation/emission wavelength of 485/520 nm. Four frames per chip at 10X magnification were selected for each treatment, and particles were counted using Fiji/ImageJ.

### Viability Assay

The cell viability was assessed using the LIVE/DEAD staining kit (Thermo Fisher Scientific). The neuronal channel of the Brain-Chips was incubated for 30 min in PBS containing 1 μM calcein AM and 2 μM ethidium homodimer-1 (EthD1). The channel was then washed with PBS and imaged under a motorized fluorescent microscope (Zeiss confocal microscope). Four frames per chip at 10X magnification were selected for each treatment, and particles were counted using Fiji/ImageJ. Data were expressed as the average live cells/total number of cells (sum of calcein-AM positive and ethidium homodimer positive). In order to confirm the efficiency and reliability of this assay, we set up a positive control (DMSO treatment) and negative control (no treatment) to do the parallel experiment with the αSyn treatment.

### Cytokine Secretion

The levels of TNF-α, and IL-6 were measured by commercial ELISA kit (Abcam) according to the manufacturers’ instructions. The assays were performed in duplicate in 96-well plates, and the results were presented as picograms per milliliter.

### RNA isolation and sequencing

RNA was extracted using TRIzol (TRI reagent, Sigma) according to manufacturer’s guidelines. The collected samples were submitted to GENEWIZ South Plainfield, NJ, for next-generation sequencing. After quality control and RNA-seq library preparation the samples were sequenced with Illumina HiSeq 2×150 system using sequencing depth ∼50M paired-end reads/sample.

### RNA sequencing bioinformatics

Using Trimmomatic v.0.36 we trimmed the sequence reads and filtered-out poor quality nucleotides and possible adapter sequences. The remained trimmed reads were mapped to the homo sapience reference genome GRCh38 using the STAR aligner v2.5.2b which generated the BAM files. Using the BAM files, we calculated for each sample the unique gene hit-counts by applying the feature Counts from the subread package v.1.5.2. Note that only unique reads that fell within the exon region were counted. Finally, the generated hit-counts were used to perform Differentially Gene Expression (DGE) analysis using the “DESeq2” R package by Bioconductor.

### GO term enrichment analysis

The gene sets resulted from the DGE analyses were subjected to Gene Ontology (GO) enrichment analysis. The GO terms enrichment analysis was performed using the Gene Ontology knowledgebase (Gene Ontology Resource http://geneontology.org/).

### GTEx human adult substantia nigra samples selection procedure

GTEx Portal provides 114 RNA-seq samples for human adult substantia nigra. 8 representative samples out of the 114 samples were selected and combined with our 8 samples from our Brain-Chip and conventional cell culture samples to generate a balanced dataset. For the selection of the 8 representative samples we used the following criteria: (1) The samples belonged to donors who were reasonably healthy and they had fast and unexpected deaths from natural causes; and (2) Have the smaller transcriptomic distances^83^ from the average transcriptomic expression profile of the samples that satisfy criterion (1). Next, we used the “removeBatchEffect” function of the “limma” R package in order to remove shifts in the means between our samples (Brain-Chip and conventional cell cultures) and the 8 human substantia nigra samples that we retrieved from GTEx portal. The dataset was used to perform DGE analyses between the different conditions. For the DGE analyses, we used the ‘DESeq2’ R package by bioconductor^84^.

### Statistical Analysis

All experiments were performed with controls (monomers or PBS) side-by-side and in random order and they were reproduced for at least two times to confirm data reliability. GraphPad Prism was used to perform statistical analyses (GraphPad Software). All numeric results are shown as mean ± standard error of the mean (SEM) and represent data from a minimum of two independent experiments of distinct sample measurements (n>3). Analysis of significance was performed by using two-way ANOVA with Tukey’s multiple comparisons test or unpaired t-test depending on the data sets. Significant differences are depicted as follows: *\*P*<0.05, \*\**P*<0.01, *\*\*\*P*< 0.001, and \*\*\*\**P<*0.0001.

## References

1. Peng, C., Gathagan, R. J. & Lee, V. M. Y. Distinct α-Synuclein strains and implications for heterogeneity among α-Synucleinopathies. Neurobiology of Disease vol. 109 209–218 (2018).

2. Peelaerts, W., Bousset, L., Baekelandt, V. & Melki, R. ?-Synuclein strains and seeding in Parkinson’s disease, incidental Lewy body disease, dementia with Lewy bodies and multiple system atrophy: similarities and differences. Cell and Tissue Research vol. 373 195–212 (2018).

3. Luk, K. C. et al. Pathological α-synuclein transmission initiates Parkinson-like neurodegeneration in nontransgenic mice. Science 338, 949–53 (2012).

4. Goedert, M., Jakes, R. & Spillantini, M. G. The Synucleinopathies: Twenty Years on. Journal of Parkinson’s Disease vol. 7 S53–S71 (2017).

5. Ouzounoglou, E. et al. In silico modeling of the effects of alpha-synuclein oligomerization on dopaminergic neuronal homeostasis. BMC Syst. Biol. 8, (2014).

6. El Agnaf, O. M. A. et al. Detection of oligomeric forms of α synuclein protein in human plasma as a potential biomarker for Parkinson’s disease. FASEB J. 20, 419–425 (2006).

7. Tokuda, T. et al. Detection of elevated levels of α-synuclein oligomers in CSF from patients with Parkinson disease. Neurology 75, 1766–1772 (2010).

8. Beach, T. G. et al. Multi-organ distribution of phosphorylated α-synuclein histopathology in subjects with Lewy body disorders. Acta Neuropathol. 119, 689–702 (2010).

9. Donadio, V. et al. Skin nerve a-synuclein deposits A biomarker for idiopathic Parkinson disease. Neurology 82, 1362–1369 (2014).

10. Kortekaas, R. et al. Blood-brain barrier dysfunction in Parkinsonian midbrain in vivo. Ann. Neurol. 57, 176–179 (2005).

11. Rektor, I. et al. Impairment of brain vessels may contribute to mortality in patients with Parkinson’s disease. Mov. Disord. 27, 1169–1172 (2012).

12. Sui, Y.-T., Bullock, K. M., Erickson, M. A., Zhang, J. & Banks, W. A. Alpha synuclein is transported into and out of the brain by the blood-brain barrier. Peptides 62, 197–202 (2014).

13. Lee, H. & Pienaar, I. S. Disruption of the blood-brain barrier in parkinson’s disease: Curse or route to a cure? Front. Biosci. -Landmark 19, 272–280 (2014).

14. Carvey, P. M. et al. 6-Hydroxydopamine-induced alterations in blood-brain barrier permeability. Eur. J. Neurosci. 22, 1158–1168 (2005).

15. Peelaerts, W. et al. α-Synuclein strains cause distinct synucleinopathies after local and systemic administration. Nature 522, 340–344 (2015).

16. Stefanis, L., Larsen, K. E., Rideout, H. J., Sulzer, D. & Greene, L. A. Expression of A53T mutant but not wild-type α-synuclein in PC12 cells induces alterations of the ubiquitin-dependent degradation system, loss of dopamine release, and autophagic cell death. J. Neurosci. 21, 9549–9560 (2001).

17. Vekrellis K., Xilouri M., Emmanouilidou E., S. L. Inducible over-expression of wild type alpha-synuclein in human neuronal cells leads to caspase-dependent non-apoptotic death. J. Neurochem. 109, 1348–1362 (2009).

18. Kuan, W.-L. et al. α-Synuclein pre-formed fibrils impair tight junction protein expression without affecting cerebral endothelial cell function. Exp. Neurol. 285, 72–81 (2016).

19. Lane, E. & Dunnett, S. Animal models of Parkinson’s disease and L-dopa induced dyskinesia: How close are we to the clinic? Psychopharmacology vol. 199 303–312 (2008).

20. Banks, W. A., Kovac, A. & Morofuji, Y. Neurovascular unit crosstalk: Pericytes and astrocytes modify cytokine secretion patterns of brain endothelial cells. J. Cereb. Blood Flow Metab. 38, 1104–1118 (2018).

21. Kaisar, M. A. et al. New experimental models of the blood-brain barrier for CNS drug discovery. Expert Opinion on Drug Discovery vol. 12 89–103 (2017).

22. Bhatia, S. N. & Ingber, D. E. Microfluidic organs-on-chips. Nature Biotechnology vol. 32 760–772 (2014).

23. Haring, A. P., Sontheimer, H. & Johnson, B. N. Microphysiological Human Brain and Neural Systems-on-a-Chip: Potential Alternatives to Small Animal Models and Emerging Platforms for Drug Discovery and Personalized Medicine. Stem Cell Rev. Reports 13, 381–406 (2017).

24. Kasendra, M. et al. Duodenum Intestine-Chip for preclinical drug assessment in a human relevant model. Elife 9, (2020).

25. Huh, D. et al. A human disease model of drug toxicity-induced pulmonary edema in a lung-on-a-chip microdevice. Sci. Transl. Med. 4, (2012).

26. Jang, K. J. et al. Reproducing human and cross-species drug toxicities using a Liver-Chip. Sci. Transl. Med. 11, (2019).

27. Agarwal, A., Goss, J. A., Cho, A., McCain, M. L. & Parker, K. K. Microfluidic heart on a chip for higher throughput pharmacological studies. Lab Chip 13, 3599–3608 (2013).

28. Moreno, E. L. et al. Differentiation of neuroepithelial stem cells into functional dopaminergic neurons in 3D microfluidic cell culture. Lab Chip 15, 2419–2428 (2015).

29. Freundt, E. C. et al. Neuron-to-neuron transmission of α-synuclein fibrils through axonal transport. Ann. Neurol. 72, 517–524 (2012).

30. Vatine, G. D. et al. Human iPSC-Derived Blood-Brain Barrier Chips Enable Disease Modeling and Personalized Medicine Applications. Cell Stem Cell 24, 995-1005.e6 (2019).

31. Park, T. E. et al. Hypoxia-enhanced Blood-Brain Barrier Chip recapitulates human barrier function and shuttling of drugs and antibodies. Nat. Commun. 10, (2019).

32. Shin, Y. et al. Blood-Brain Barrier Dysfunction in a 3D In Vitro Model of Alzheimer’s Disease. Adv. Sci. (Weinheim, Baden-Wurttemberg, Ger. 6, 1900962 (2019).

33. Ahn, S. I. et al. Microengineered human blood–brain barrier platform for understanding nanoparticle transport mechanisms. Nat. Commun. 11, (2020).

34. Choi, J. H., Santhosh, M. & Choi, J. W. In vitro blood-brain barrier-integrated neurological disorder models using a microfluidic device. Micromachines vol. 11 (2020).

35. Ganjam, G. K. et al. Mitochondrial damage by α-synuclein causes cell death in human dopaminergic neurons. Cell Death Dis. 10, 865 (2019).

36. Hirsch, E. C., Vyas, S. & Hunot, S. Neuroinflammation in Parkinson’s disease. Park. Relat. Disord. 18, (2012).

37. Obermeier, B., Daneman, R. & Ransohoff, R. M. Development, maintenance and disruption of the blood-brain barrier. Nature Medicine vol. 19 1584–1596 (2013).

38. Qian, T. et al. Directed differentiation of human pluripotent stem cells to blood-brain barrier endothelial cells. Sci. Adv. 3, e1701679 (2017).

39. Kniesel, U. & Wolburg, H. Tight junctions of the blood-brain barrier. Cell. Mol. Neurobiol. 20, 57–76 (2000).

40. Shi, L., Zeng, M., Sun, Y. & Fu, B. M. Quantification of blood-brain barrier solute permeability and brain transport by multiphoton microscopy. J. Biomech. Eng. 136, (2014).

41. Yuan, W., Lv, Y., Zeng, M. & Fu, B. M. Non-invasive measurement of solute permeability in cerebral microvessels of the rat. Microvasc. Res. 77, 166–173 (2009).

42. Battle A, Brown C D, Engelhardt B E, M. S. B. Genetic effects on gene expression across human tissues. Nature 550, 204–213 (2017).

43. Hesari, Z. et al. A hybrid microfluidic system for regulation of neural differentiation in induced pluripotent stem cells. J. Biomed. Mater. Res. -Part A 104, 1534–1543 (2016).

44. Samal, P., van Blitterswijk, C., Truckenmüller, R. & Giselbrecht, S. Grow with the Flow: When Morphogenesis Meets Microfluidics. Adv. Mater. 31, (2019).

45. Sances, S. et al. Human iPSC-Derived Endothelial Cells and Microengineered Organ-Chip Enhance Neuronal Development. Stem Cell Reports 10, 1222–1236 (2018).

46. Marques, O. & Outeiro, T. F. Alpha-synuclein: From secretion to dysfunction and death. Cell Death and Disease vol. 3 (2012).

47. Volpicelli-Daley, L. A. et al. Exogenous α-Synuclein Fibrils Induce Lewy Body Pathology Leading to Synaptic Dysfunction and Neuron Death. Neuron 72, 57–71 (2011).

48. Anderson, J. P. et al. Phosphorylation of Ser-129 is the dominant pathological modification of α-synuclein in familial and sporadic lewy body disease. J. Biol. Chem. 281, 29739–29752 (2006).

49. Arawaka, S., Sato, H., Sasaki, A., Koyama, S. & Kato, T. Mechanisms underlying extensive Ser129-phosphorylation in α-synuclein aggregates. Acta Neuropathol. Commun. 5, 48 (2017).

50. Moon HE, P. S. Mitochondrial Dysfunction in Parkinson’s Disease. Exp. Neurobiol. 24, 103–116 (2015).

51. Sivandzade, F., Bhalerao, A. & Cucullo, L. Analysis of the Mitochondrial Membrane Potential Using the Cationic JC-1 Dye as a Sensitive Fluorescent Probe. BIO-PROTOCOL 9, (2019).

52. Gelders, G., Baekelandt, V. & Van der Perren, A. Linking Neuroinflammation and Neurodegeneration in Parkinson’s Disease. J. Immunol. Res. 2018, 4784268 (2018).

53. Desai, B. S., Monahan, A. J., Carvey, P. M. & Hendey, B. Blood-brain barrier pathology in Alzheimer’s and Parkinson’s disease: Implications for drug therapy. in Cell Transplantation vol. 16 285–299 (Cognizant Communication Corporation, 2007).

54. Munji, R. N. et al. Profiling the mouse brain endothelial transcriptome in health and disease models reveals a core blood–brain barrier dysfunction module. Nat. Neurosci. 22, 1892–1902 (2019).

55. Gandhi, P. N., Chen, S. G. & Wilson-Delfosse, A. L. Leucine-rich repeat kinase 2 (LRRK2): A key player in the pathogenesis of Parkinson’s disease. Journal of Neuroscience Research vol. 87 1283–1295 (2009).

56. Wakabayashi, K. et al. Synphilin-1 is present in lewy bodies in Parkinson’s disease. Ann. Neurol. 47, 521–523 (2000).

57. Youdim, M. B. H. & Bakhle, Y. S. Monoamine oxidase: Isoforms and inhibitors in Parkinson’s disease and depressive illness. British Journal of Pharmacology vol. 147 (2006).

58. Carpanini, S. M., Torvell, M. & Morgan, B. P. Therapeutic inhibition of the complement system in diseases of the central nervous system. Frontiers in Immunology vol. 10 (2019).

59. Del Giudice, R. et al. Amyloidogenic variant of apolipoprotein A-I elicits cellular stress by attenuating the protective activity of angiogenin. Cell Death Dis. 5, (2014).

60. Gosselet, F. et al. Transcriptional profiles of receptors and transporters involved in brain cholesterol homeostasis at the blood-brain barrier: Use of an in vitro model. Brain Res. 1249, 34–42 (2009).

61. Furuno, T. et al. Expression polymorphism of the blood-brain barrier component P-glycoprotein (MDR1) in relation to Parkinson’s disease. Pharmacogenetics 12, 529–534 (2002).

62. Jin, U., Park, S. J. & Park, S. M. Cholesterol metabolism in the brain and its association with Parkinson’s disease. Experimental Neurobiology vol. 28 554–567 (2019).

63. Yoon, Y.-S. et al. Is trehalose an autophagic inducer? Unraveling the roles of non-reducing disaccharides on autophagic flux and alpha-synuclein aggregation. Cell Death Dis. 8, e3091 (2017).

64. Hoffmann, A.-C. et al. Extracellular aggregated alpha synuclein primarily triggers lysosomal dysfunction in neural cells prevented by trehalose. Sci. Rep. 9, 544 (2019).

65. Larocca, T. J. et al. Translational evidence that impaired autophagy contributes to arterial ageing. J. Physiol. 590, 3305–3316 (2012).

66. Rodríguez-Navarro, J. A. et al. Trehalose ameliorates dopaminergic and tau pathology in parkin deleted/tau overexpressing mice through autophagy activation. Neurobiol. Dis. 39, 423–38 (2010).

67. Lan, D. M. et al. Effect of trehalose on PC12 cells overexpressing wild-type or A53T mutant α-synuclein. Neurochem. Res. 37, 2025–2032 (2012).

68. Lázaro, D. F., Pavlou, M. A. S. & Outeiro, T. F. Cellular models as tools for the study of the role of alpha-synuclein in Parkinson’s disease. Experimental Neurology vol. 298 162–171 (2017).

69. Wood, S. J. et al. α-Synuclein fibrillogenesis is nucleation-dependent: Implications for the pathogenesis of Parkinson’s disease. J. Biol. Chem. 274, 19509–19512 (1999).

70. McGeer, P. L., Itagaki, S., Boyes, B. E. & McGeer, E. G. Reactive microglia are positive for HLA-DR in the: Substantia nigra of Parkinson’s and Alzheimer’s disease brains. Neurology 38, 1285–1291 (1988).

71. Lofrumento, D. D. et al. MPTP-induced neuroinflammation increases the expression of pro-inflammatory cytokines and their receptors in mouse brain. Neuroimmunomodulation 18, 79–88 (2010).

72. Mogi, M. et al. Tumor necrosis factor-α (TNF-α) increases both in the brain and in the cerebrospinal fluid from parkinsonian patients. Neurosci. Lett. 165, 208–210 (1994).

73. Mogi, M., Harada, M., Kondo, T., Riederer, P., Inagaki, H., Minami, M. and Nagatsu, T. Interleukin-1β, interleukin-6, epidermal growth factor and transforming growth factor-α are elevated in the brain from parkinsonian patients. Neurosci. Lett. 180, 147–150 (1994).

74. Lee, H. J., Suk, J. E., Bae, E. J. & Lee, S. J. Clearance and deposition of extracellular α-synuclein aggregates in microglia. Biochem. Biophys. Res. Commun. 372, 423–428 (2008).

75. Man, S. et al. CXCL12-induced monocyte-endothelial interactions promote lymphocyte transmigration across an in vitro blood-brain barrier. Sci. Transl. Med. 4, (2012).

76. Zhao, C., Ling, Z., Newman, M. B., Bhatia, A. & Carvey, P. M. TNF-α knockout and minocycline treatment attenuates blood-brain barrier leakage in MPTP-treated mice. Neurobiol. Dis. 26, 36–46 (2007).

77. Jangula, A. & Murphy, E. J. Lipopolysaccharide-induced blood brain barrier permeability is enhanced by alpha-synuclein expression. Neurosci. Lett. 551, 23–27 (2013).

78. Gray, M. T. & Woulfe, J. M. Striatal blood-brain barrier permeability in Parkinson’s disease. J. Cereb. Blood Flow Metab. 35, 747–750 (2015).

79. Rite, I., Machado, A., Cano, J. & Venero, J. L. Blood-brain barrier disruption induces in vivo degeneration of nigral dopaminergic neurons. J. Neurochem. 101, 1567–1582 (2007).

80. Logsdon, A. F., Erickson, M. A., Rhea, E. M., Salameh, T. S. & Banks, W. A. Gut reactions: How the blood-brain barrier connects the microbiome and the brain. Exp. Biol. Med. (Maywood). 243, 159–165 (2018).

81. Gonçalves, A., Ambrósio, A. F. & Fernandes, R. Regulation of claudins in blood-tissue barriers under physiological and pathological states. Tissue Barriers 1, e24782 (2013).

82. Maoz, B. M. et al. A linked organ-on-chip model of the human neurovascular unit reveals the metabolic coupling of endothelial and neuronal cells. Nat. Biotechnol. 36, 865–877 (2018).

83. Manatakis, D. V., VanDevender, A. & Manolakos, E. S. An information-theoretic approach for measuring the distance of organ tissue samples using their transcriptomic signatures. Bioinformatics. btaa654, https://doi.org/10.1093/bioinformatics/btaa654 (2020)

84. Love, M. I., Huber, W. & Anders, S. Moderated estimation of fold change and dispersion for RNA-seq data with DESeq2. Genome Biol. 15, (2014).

